# Deep quantitative phosphoproteomics identifies non-canonical pH-sensitive yeast phosphorylation networks

**DOI:** 10.64898/2026.04.13.718172

**Authors:** Xinya Su, Aarushi Gajri, Matthew Torres

**Affiliations:** School of Biological Sciences, Georgia Institute of Technology, Atlanta, GΑ 30033

## Abstract

Changes in cellular pH act as potent upstream signals that rewire phosphorylation networks controlling cellular function and disease. In yeast, this response is thought to arise mainly from pH-dependent perturbations to membrane integrity and cell wall stress, which are relayed through TORC2- and PKC-dependent signaling pathways. Yet several proteins exhibit acid-dependent phosphorylation independently of TORC2 and PKC, pointing to additional, uncharacterized acid stress signaling mechanisms. To investigate this possibility, we performed SILAC-based deep phosphoproteomics to distinguish phosphoproteome changes that are dependent on plasma membrane and cell wall stress from those that are independent. Across more than 19,000 unique phosphosites, we identified over 1,000 sites whose acid stress–dependent regulation occurs outside the canonical acid stress pathways. These noncanonical targets were significantly enriched in proteins associated with the plasma membrane, GTPase-mediated signaling, and endocytosis. Motif analysis revealed enrichment of acidophilic substrates implicating the membrane-tethered casein kinase Yck1 as a major mediator of this response. In contrast, canonical acid stress signaling preferentially involved basophilic kinase substrates, while acid-repressed responses were enriched for proline-containing phosphosites involved in cell cycle progression. Collectively, these results uncover a distinct acid-responsive phosphorylation network that operates independently of plasma membrane and cell wall integrity signaling.

## INTRODUCTION

Protein structure and function are inherently sensitive to pH and intracellular pH (pHi) homeostasis is therefore essential for proteome function and cell survival (1–4). Dysregulation of pHi contributes to dynamic functionality of the proteome in cancer and stem cells, with multiple examples of pHi having discrete effects on specific signaling pathways and proteins (5–8). Several proteins are directly sensitive to intracellular proton concentration, and in these cases the proton itself can function as a second messenger that activates or modulates signaling (9–13). Most examples have been found to occur at the plasma membrane or other intracellular membranes. These observations provide supporting evidence for the existence of proteome networks, and possibly pathways, that are not only sensitive to the indirect consequences of dynamic pHi but also regulated by it directly.

Significant changes in protein phosphorylation are concomitant with acute pH stress (14–17). In most cases, the mechanism linking pH changes to phosphorylation is unknown and where it has been defined, the available evidence more often supports indirect, secondary effects than direct proton-mediated activation. Acid stress, for example, is known to influence membrane physical properties including global deformation and local dynamic membrane remodeling through acid/base chemistry of membrane lipids (18). In yeast, two canonical pH-response pathways are thought to feed from this pH-to-membrane deformation mechanism: a plasma membrane (PM) stress pathway mediated by TORC2 and eisosome-associated factors that lead to activation of Ypk1 kinase and other downstream proteins (19–21), as well as the cell wall integrity (CWI) pathway mediated by the Wsc-family mechanosensing sensors and Mid2 that promote activation of Rom2, Rho1, Pkc1, Bck1, and Slt2 kinases (22).

Despite their observed connection to pH-coupled membrane stress, PM stress and CWI pathways mediated by TORC2 and PKC do not explain the entirety of early phosphorylation responses to acid stress, suggesting that additional pH response pathways exist (15). To evaluate this hypothesis systematically, we conducted a multi-dimensional quantitative phosphoproteomics experiment that decouples the yeast phosphoproteome response to acetic acid stress in the presence or absence of a PM/CWI signaling blockade.

## RESULTS

### Quantitative deep phosphoproteomics distinguishes intracellular acidification from acid-induced cell wall/plasma membrane stress

Our goal was to distinguish acid stress-activated phosphorylations that are either dependent or independent on canonical TORC/PKC response pathways that are dependent on plasma membrane stress. To do this, we designed a quantitative deep-phosphoproteomics experiment wherein stable isotope labels were used to distinguish three different cell treatments: Untreated (Light; L), acid-treated (Medium; M), and sorbitol-pre-equilibrated/acid-treated (Heavy; H) (**Fig. 1A**). To avoid bias due to changes in protein expression, quantitative phosphopeptide data were also normalized by changes in protein abundance gathered from a whole proteome analysis of the same samples carried out in parallel. Previous studies have shown that acid stress activates both TORC2-Ypk1 and PKC/CWI signaling in yeast, and that osmotic support with sorbitol can suppress or attenuate pH-associated cell-surface stress responses, consistent with a role for plasma membrane and cell wall stabilization in blunting these pathways (17, 23–25). Accordingly, sites whose phosphorylation increases under acid stress alone but not in sorbitol-equilibrated cells represent canonical TORC2/PKC/acid-dependent phosphorylations. Phosphosites that are unaffected by the presence or absence of sorbitol represent non-canonical acid stress phosphorylation that is the primary focus of our work.

**Fig. 1.**
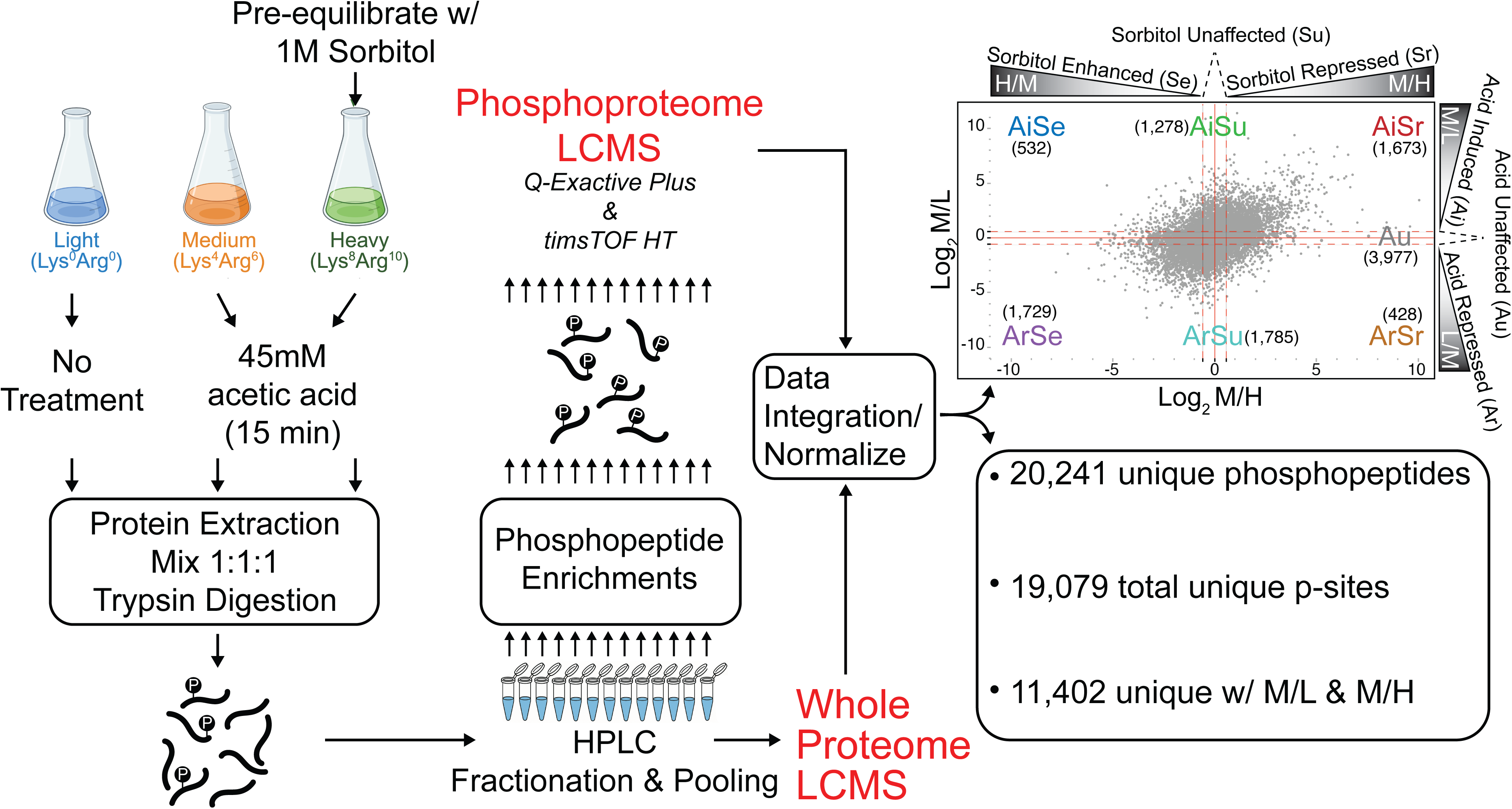
Experimental workflow for deep phosphoproteomic analysis of the yeast acid stress response in the presence and absence of sorbitol. Triple SILAC labeling of yeast cells undergoing no treatment, acid stress alone or acid stress in cells equilibrated to 1M sorbitol to block acid-dependent TORC2/CWI signaling. After hi-pH reverse phase LC fractionation and re-pooling of peptides, each of 13 fractions were analyzed by LCMS to establish protein abundance profiles across the three conditions. Phosphopeptides from each fraction were enriched and analyzed separately by LCMS to quantify phosphopeptide abundances. Whole proteome and phosphoproteome data were then integrated and normalized to account for changes in protein abundance that reflect accurate changes in phosphorylation state change, which were divided into 7 distinct classes depending on the acid response alone (Log_2_M/L) versus the acid response in sorbitol-equilibrated cells (Log_2_M/H).

In our experiment, the effect of acid alone is given by the M/L ratio, wherein a ratio >1 ([Log_2_]_M/L_ >0) indicates an increase in phosphorylation upon acid treatment (acid induced, Ai) while acid-unaffected (Au) sites correspond to a ratio near 1 ([Log_2_]_M/L_ ∼0). Of these acid-induced sites, those that are unaffected by sorbitol are distinguished by the M/H ratio, wherein ratios >1 ([Log_2_]_M/H_ >0) indicate sorbitol repression of the acid responsive phosphorylation (Sr) whereas those with M/H ratios near 1 ([Log_2_]_M/H_ ∼0) correspond with sorbitol unaffected sites (Su). Together, a total of six response classes were used to analyze the data: Acid induced/Sorbitol enhanced (AiSe; Log_2_M/L>0.585, Log_2_M/H<-0.585); Acid induced/Sorbitol unaffected (AiSu; Log_2_M/L>0.585, -0.585<Log_2_M/H<0.585); Acid induced/Sorbitol repressed (AiSr; Log_2_M/L>0.585, Log_2_M/L>0.585); Acid repressed/Sorbitol enhanced (ArSe; Log_2_M/L<0.585, Log_2_M/H<-0.585); Acid repressed/Sorbitol unaffected (ArSu, Log_2_M/L<0.585, -0.585<Log_2_M/H<0.585); and Acid repressed/Sorbitol repressed (ArSr; Log_2_M/L<0.585, Log_2_M/L>0.585). Overall, the experiment resulted in detection of 19,079 unique phosphorylation sites covering >70% of the known yeast phosphoproteome (*supplemental information*). Of these, 11,402 sites in M/L or M/H could be compared across all three conditions, 5,951 of which show a significant change in conditional response across both conditional comparisons (M/H or M/L) (p<0.05) (**Supplementary Fig. S1A,B**). Comparative analysis of biological processes, cellular compartments and molecular functions revealed pronounced and significant enrichment of TOR complex and several TORC2-associated bio processes in the Sr relative to Se response classes, as expected for phosphorylations that depend on acid-mediated membrane disruption (**Fig. 2A,B**). Acid-induced phosphorylations were similarly enriched in actin-based processes but even more so in ribonucleoprotein granules and endocytosis, while acid repressed phosphorylations were comparatively enriched in mitotic cell cycle process, chromatin, and chromatin remodeling (**Fig. 2C**). Finally, a more precise comparison between AiSu and AiSr response classes, which represents the primary focal point of this work, showed strong relative enrichment for GTPases, signal transduction, endocytosis, and intracellular vesicles that provided the first glimpse into a target class for TORC2/PKC-independent acid stress targets (**Fig. 2D**). A more refined analysis of the AiSu/AiSr comparison reinforced this finding, showing prevalent enrichment of plasma membrane-associated terms involving signal transduction, cell communication and vesicle-mediated transport (**Supplemental Fig. S2**).

**Fig. 2.**
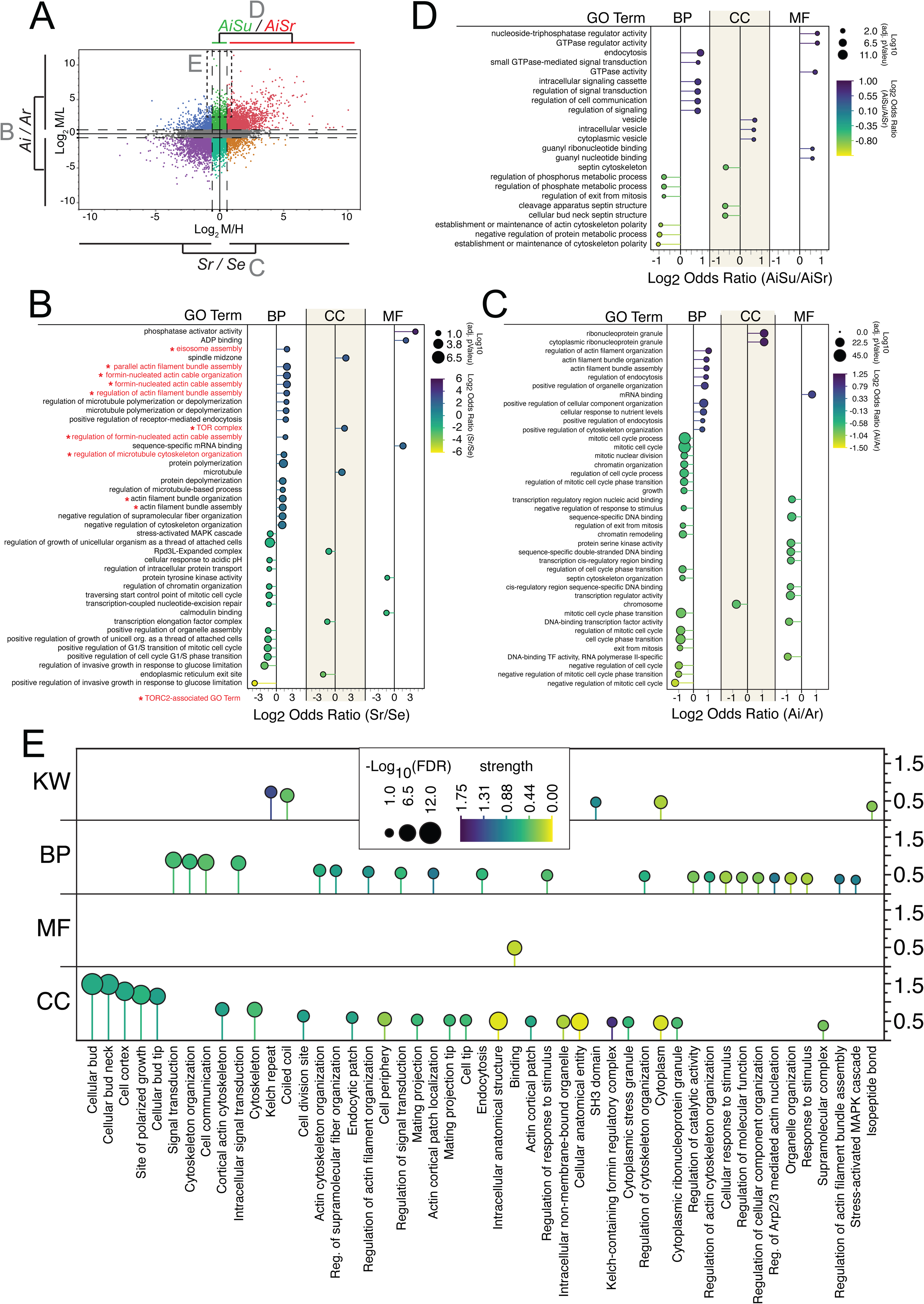
Gene ontology enrichment of the major acid/sorbitol response classes. (A) Log2 M/H vs. M/L plot for phosphopeptides observed in the experiment. Gene ontology enrichment comparisons discussed in later panels is indicated along with the panel letter. Multi-comparison gene ontology enrichments produced using g:Profiler were converted into log odds values and then ratioed between the indicated response classes in panels B-E. (B) Relative comparison in gene ontology enrichment between sorbitol induced (Si) and sorbitol repressed (Sr) responses. (C) Relative comparison in gene ontology enrichment between acid induced (Ai) and acid repressed (Ar) responses. (D) Relative comparison in gene ontology enrichment between acid induced/sorbitol unaffected (AiSu) and acid induced/sorbitol repressed (AiSr) responses. (E) Gene ontology enrichment in proteins identified as having an extreme acid response and minimal sorbitol response (Log_2_M/L>2.5, -1<Log_2_M/H<1). BP, Biological Process; CC, Cellular Component; MF, Molecular Function; KW, Kegg WikiPathways. All data shown is filtered at adjusted p<0.05.

Lastly, we analyzed the enrichment of gene ontologies for proteins that undergo extreme phosphorylation in acetic acid and minimal effect by sorbitol (**Fig. 2A** dashed box). These proteins were highly enriched in plasma membrane localized functions at the bud neck or sites of polarized growth involving signal transduction, cell communication and cytoskeletal organization (**Fig. 2E**). Interestingly, proteins containing kelch repeats, which are often involved in protein interactions, signal transduction, ubiquitin-mediated degradation, and membrane trafficking, were extremely significantly enriched (q=0.001) with 5 out of 7 total ontology members observed (26, 27). Taken together, these data reveal a total 1,767 phosphosites in the AiSu response class and provide support of the hypothesis that acid stress promotes TORC2/PKC-independent signaling centered on plasma membrane localized proteins involved in signal transduction, cytoskeletal organization, and endocytosis.

### Quantitative MS distinguishes sorbitol-repressed TORC2/PKC signaling from sorbitol-insensitive acid stress phosphorylation

The design of our MS experiment was predicated on previous evidence that acid-dependent activation of TORC2 and PKC pathways can be repressed by osmo-protectants, while others cannot. Ypk1-S644/S662 are direct TORC2 substrate sites undergoing acid-dependent phosphorylation that is repressed in the presence of sorbitol (23) (**Fig. 3A**), an observation we confirmed by quantitative immunoblotting (**Fig. 3B,C**). This observation is further supported by immunoblot evidence for Slt2, a downstream substrate of PKC in the cell wall integrity pathway (**Fig. 3A**), which is activated by acid stress but repressed by sorbitol (**Supplemental Fig. S3**). In contrast to Ypk1 and Slt2, we discovered that phosphorylation of the membrane-associated G protein Gγ subunit, Ste18-S3, which undergoes robust acid stress phosphorylation in its N-terminal disordered tail (15), is unaffected by sorbitol (**Fig. 3A-C**).

**Fig. 3.**
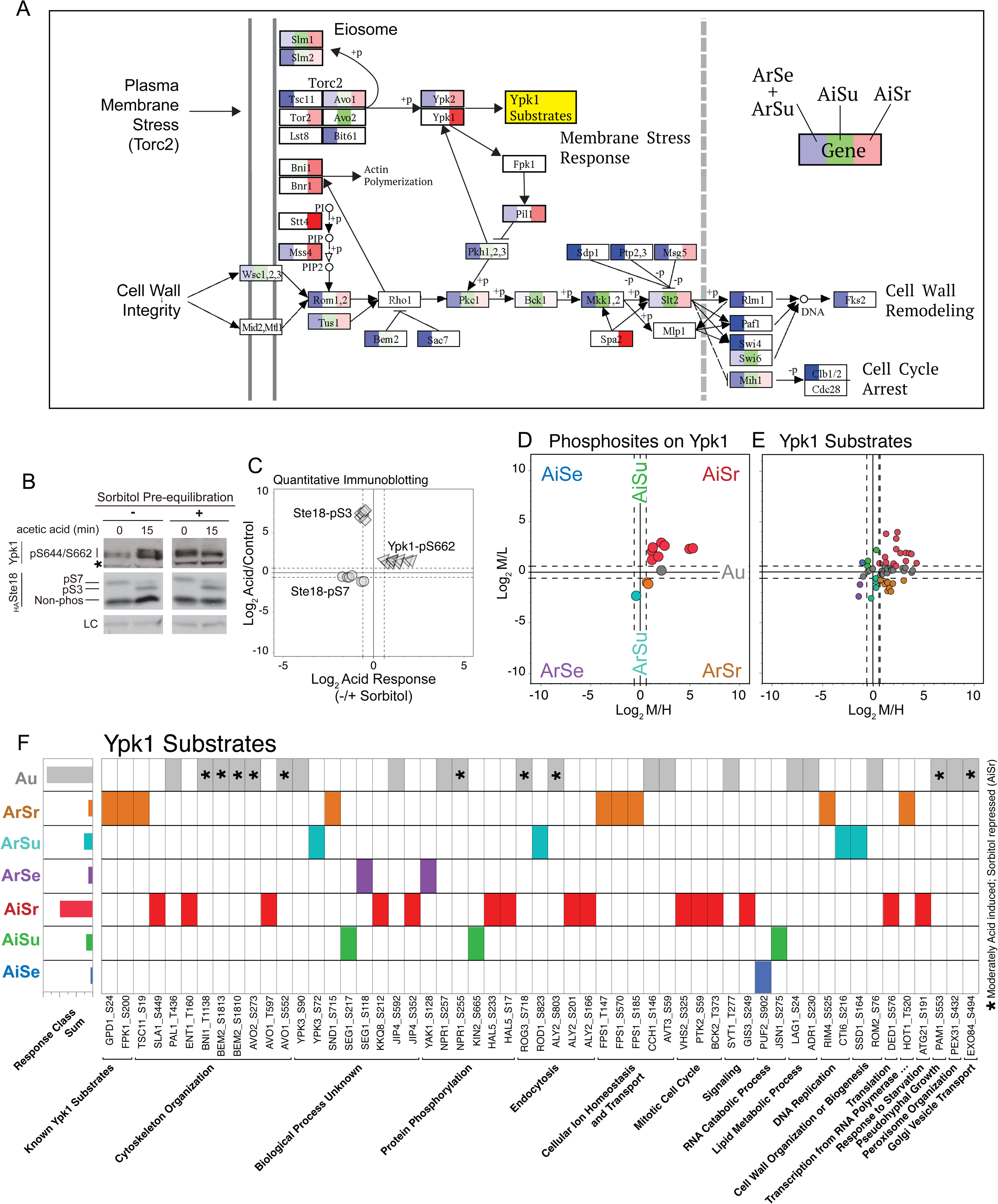
TORC2 and CWI pathways are induced by acid stress but not when cells are pre-equilibrated in sorbitol. (A) Customized Kegg pathway map highlighting the number of phosphosites observed in the ArSe+ArSu, AiSu, or AiSr response classes relative to proteins in the TORC2 and CWI pathways. Torc2, Ypk1, and Ypk1 substrates (yellow) are primary TORC2-responsive proteins expected to be enriched in the AiSr response class. (B) Quantitative immunoblotting demonstrates the site-specific acid and acid/sorbitol phosphorylation effects of Ypk1-pS662, Ste18-pS3, and Ste18-pS7. (C) Data from B were quantified across multiple replicates and plotted with respect to their Log_2_ acid/no treatment responses (y-axis) versus the Log_2_ acid response with or without sorbitol pre-equilibration (x axis). (D) Acid and acid/sorbitol effects on Ypk1 phosphosites extracted from this phosphoproteome dataset. (E) Acid and acid/sorbitol effects on phosphosites within bona fide Ypk1 substrates. (F) Phosphorylation response of sites known or predicted Ypk1 phosphosites in the yeast proteome (see Muir et al., 2014 for original source list). *Phosphosite is near the border of the AiSr response class.

We next looked for similar evidence that Ypk1 substrates and direct phosphosites are upregulated in the AiSr response class of our LCMS dataset. To do this we mapped experimentally-validated kinase/substrate and kinase/pSite pairs using data from BioGrid as well as the published literature (24). Over 70% of bona-fide Ypk1 phosphosites were enriched in the AiSr response class (**Fig. 3D**), consistent with TORC2 subunits and other known TORC2 substrates (**Fig. 3A**). At the substrate level, the vast majority of known Ypk1 substrates harbored phosphosites were also found in the AiSr response class or very adjacent Au class proximal to AiSr (**Fig. 3E,F**). Taken together, these data demonstrate the validity of the quantitative MS approach to distinguish TORC2/PKC-dependent and independent acid stress responses.

### Acid-induced/Sorbitol-unaffected phosphorylations exhibit distinctive physicochemical, sequence, and structural properties

Given that the SILAC MS dataset provides a quantitative framework to dissect acid stress signaling networks, we next asked whether it might also distinguish kinase substrate properties that differ between the response classes. We started by searching for positional enrichment of physicochemical properties and amino acids within 15-position windows surrounding the phosphosites in each response class (**Fig. 4A-F**). One of the strongest trends was the striking under-enrichment of proline-containing sequences, particularly at the +1-position relative to the phosphosite in acid repressed classes, ArSe and ArSu (**Fig. 4D,E**). Moreover, we found that, on average, the degree of phosphorylation repression under acid stress was proportional to the number of proline residues contained within the phosphopeptide sequence, suggesting a strong anti-correlation between acidic pH and phosphorylation of these sequences (**Supplemental Fig. S4**). In stark contrast, acid induced classes were statistically enriched in hydrophobic residues at the +1 position, with very clear preference for ILFVM in AiSr (**Fig. 4B,C**). The AiSu response class was also enriched in acidic residues, with D being favored at positions -3 and -2 relative to the phosphosite (**Fig. 4B**). In contrast, AiSr phosphosites are enriched in basic residues at -3, -2 and other positions upstream of the phosphosite (**Fig. 4C**). We also observed a striking trend in phosphosites that are less affected by acid stress (Au class), which tended to be near the extreme N-termini of proteins and significantly enriched in acidic residues downstream of the phosphosite (**Supplemental Fig. S5**). This enrichment trend was centered at the intersection of M/L and M/H in the Au class (the center square), indicating that these phosphosites are generally benign with respect to acid stress.

**Fig. 4.**
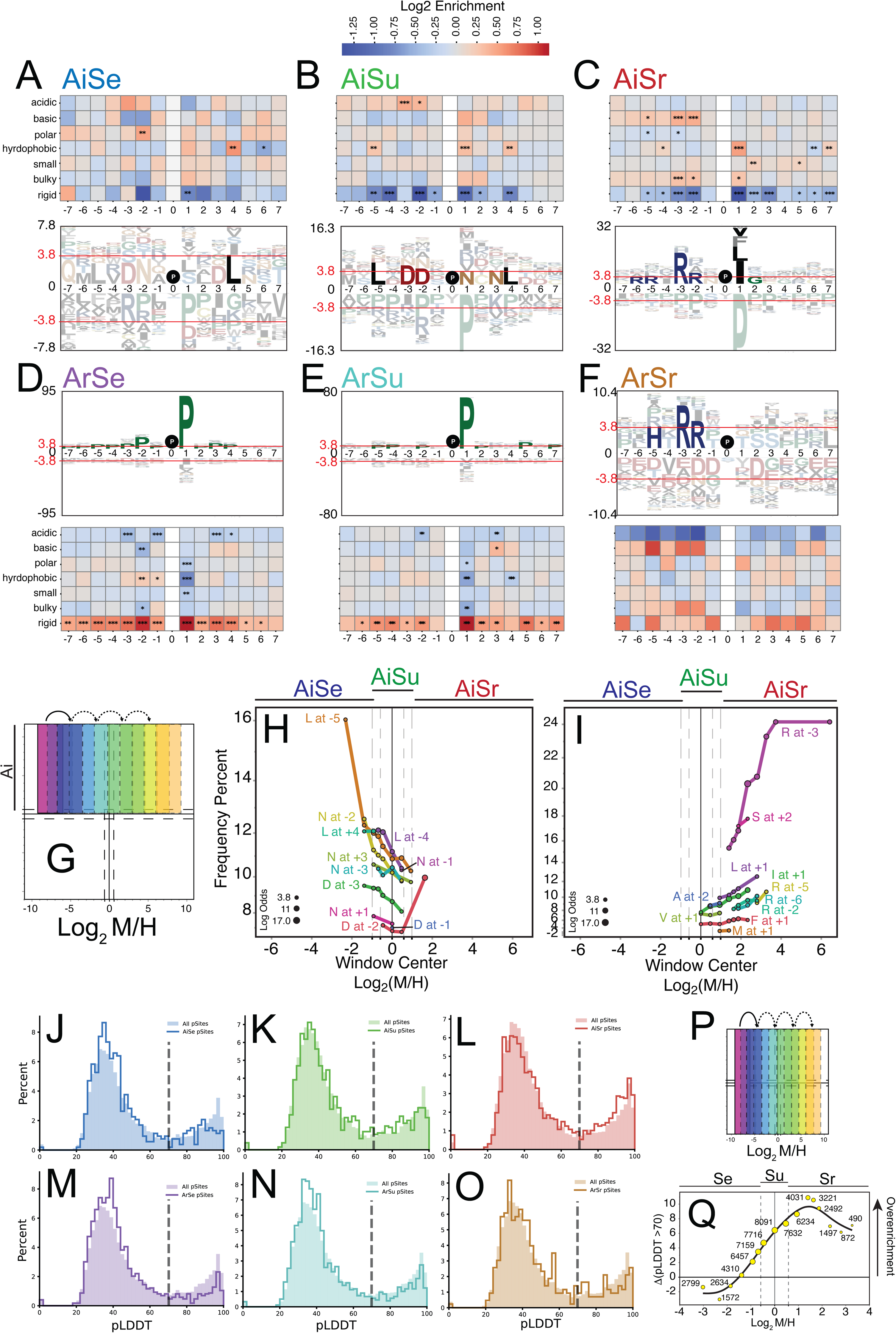
Acid induced phosphorylation occurs primarily on two distinct motifs that coincide with sorbitol repressed versus sorbitol unaffected acid responses. (A-F) Position-specific enrichment of local physicochemical properties and amino acids surrounding the acid responsive phosphoproteome. Unfaded amino acids are statistically enriched at the indicated positions relative to the phosphosite (position 0). (G) Sliding enrichment windows were used to establish trends in the enrichment of position-specific amino acids in the acid induced half of the phosphoproteomic dataset. (H,I) Enrichment trends for position-specific amino acids derived from the sliding windows. Points are plotted with respect to each window center and are divided for clarity into trend lines that move rightward starting from Log_2_M/H = 0 (I) versus those that overlap with AiSe and AiSu response classes (H). (J-O) Plot of AlphaFold 3 pLDDT score distributions in each response class (solid lines) versus the distribution of all experimental yeast phosphosites (shaded area). pLDDT scores >70 are accurately predicted fold structures (dashed grey line), while those <70 are low accuracy predictions that often coincide with less well-ordered structures. (P) Sliding enrichment windows were used to quantify an observed trend in over/under-enrichment of high-accuracy AlphaFold predictions relative to all known yeast phosphosites along the sorbitol-conditioned acid response axis. (Q) Relative difference in the pLDDT scores >70 across each of the sliding windows shown in P.

To establish whether sequence enrichment differences observed between sorbitol insensitive and sensitive acid responses were not simply due to random chance, we tracked pLogos enrichments using sliding windows along the Log_2_M/H axis (**Fig. 4G**). A clear trend emerged revealing a bifurcation in positional amino acid enrichment centered around Log_2_M/H ∼1 (**Fig. 4H,I**). To the left of this demarcation, and encompassing the AiSu response class, phosphosite sequences were enriched in leucine and asparagine both up and downstream of the phosphosite, aspartic acid at the -3 position, and asparagine at the +1 position (**Fig. 4H**). To the right of the demarcation, and encompassing the AiSr response class, phosphosite sequences were similarly enriched in +1 hydrophobic residues and prominently enriched in arginine upstream of the phosphosite (**Fig. 4I**). Thus, a clear distinction exists in the kinase substrates for canonical versus non-canonical acid stress signaling.

We also observed distinctions in the structural nature of phosphosites in the dataset. In general, most phosphosites in the yeast proteome and in our dataset exhibit AlphaFold pLDDT scores less than 70, which are classified as having low confidence for ordered structure (**Fig. 4J-O**). However, careful analysis across all dimensions of the dataset revealed a statistically enrichment of high pLDDT scores in AiSr, which was opposite to the trend found for the opposing ArSe class (**Fig. 4L,M**). To determine if this was a chance occurrence between the two response classes or an actual trend, we tracked the enrichment in pLDDT > 70 using sliding windows along the Log_2_M/H axis, demonstrating a pronounced increase in the enrichment of phosphosites in likely ordered structures (**Fig. 4P,Q**). This trend was also evident if we restricted the windowed analysis to acid induced (Log_2_M/H > 0) or acid repressed classes (Log_2_M/H < 0). These results support the hypothesis that non-canonical, TORC2/PKC-independent acid stress signaling is catalyzed by unique kinases with distinctive substrate specificities. The data also reveal a surprising correlation between the secondary structure of kinase substrates in non-canonical versus canonical acid stress signaling.

### Phosphosites with upstream histidine residues exhibit greater acid induced phosphorylation

We hypothesized that proteins exhibiting an extreme response to acid stress (in any direction) would harbor distinctive phosphosite sequences if their phosphorylated sequences happen to promote high rates of phosphorylation. To test this hypothesis, we analyzed the sequences of outlier phosphopetides falling outside the mean-centered ellipse that excluded 99% of the data (**Fig. 5A**). Significant results were only observed for outliers in the AiSr class, which revealed prominent enrichment of histidine rather than arginine at positions -4 and -3 relative to the phosphosite (**Fig. 5B**). To test whether phosphopeptides with -4H or -3H undergo greater phosphorylation in response to acid stress, we evaluated the average Log_2_M/L response for every amino acid across all positions ranging from -5 to -1. Histidine at -4 or -3, but no other amino acid or position, exhibit a significant increase in acid-dependent phosphorylation (**Fig. 5C**). Gene ontology analysis of the extreme outlier phosphopeptides in AiSr revealed TORC2 and CWI networks as well as a distinctive network anchored by the formin, Bni1, and the HECT-type E3 ligase, Rsp5, involved in actin cable assembly at the cell cortex/plasma membrane and ubiquitin-mediated endocytosis, respectively (**Fig. 5D,E**). The -4/-3H-containing phosphosites were largely accessory to these two networks, which may be due to a lack of knowledge about pH-dependent interactions in the STRING database. These results suggest that protein networks localized to the cell cortex and involving TORC2/CWI pathways, as well as cytoskeletal organization and cell polarization are primary targets of the canonical TORC2/CWI response to pH stress and that -4/-3H phosphosites are accessory to these processes.

**Fig. 5.**
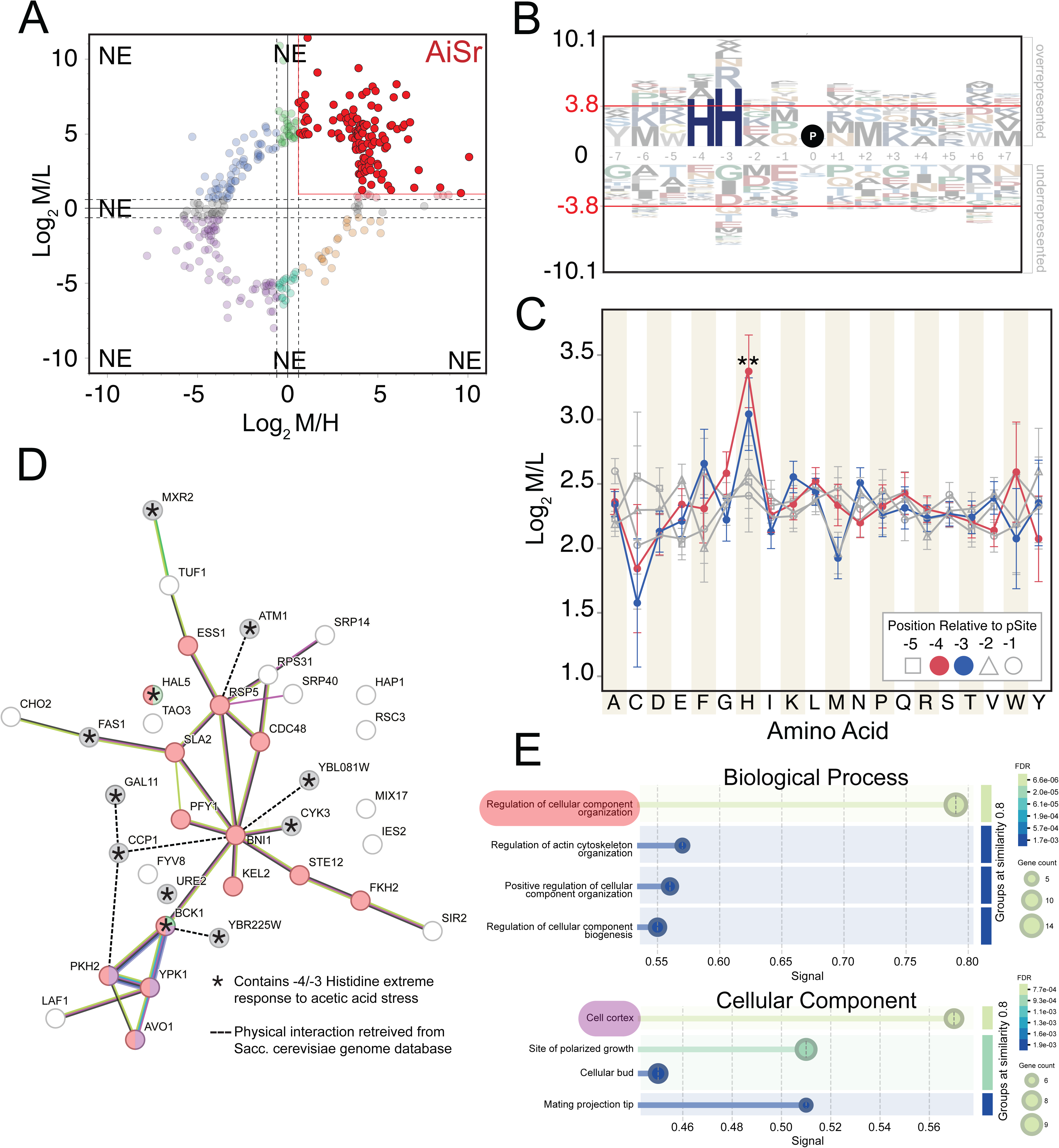
Analysis of extreme phosphorylation responses identifies histidine-neighboring phosphosites as acid hypersensitive. (A) Phosphosites outside the bounds of an ellipse covering 99% of the phosphoproteome dataset were systematically analyzed for sequence and gene ontology enrichment. NE = no significant enrichment observed. (B) pLogos plot of phosphosite-local amino acids in the AiSr extreme response class. Unfaded amino acids are significantly enriched with respect to the entire phosphoproteome dataset (p<0.05). (C) Mean acid induced response of phosphopeptides with respect to amino acid identity at positions -5 through -1 relative to the phosphosite (position 0). All AiSr phosphosites are included. (D) STRING protein network and (E) gene ontology enrichment for AiSr extreme response phosphosites. Red- and purple-colored nodes belong to ontologies shown in E. Grey and/or asterisk-containing nodes correspond to proteins harboring -4/-3H phosphosite(s).

### Enrichment of positional sequence pairs defines a motif for acid-induced, TORC2/PKC-independent phosphorylation

Sequence logos, while informative, do not report on co-enrichment across more than one position. We tested several common motif enrichment methods, which did not perform to our satisfaction, possibly due to the rich complexity of the data and the tendency for acid induced phosphosites to exist in poorly conserved disordered regions. To simplify the problem, we measured statistical enrichment of all possible pairs of amino acids across all positions surrounding the phosphosites in the data. This revealed -3/+1 and a handful of other statistically enriched positional pairs across all phosphosites (**Fig. 6A**). We selected the most prominent positional pairs and plotted the significant amino acid-pairings within these positions with respect to their enrichment in the sliding windows of the acid induced subset of the data (*refer to fig. 4G*). This revealed clear trends in the persistence of certain amino acid pairs distributed along the Log_2_M/H axis (**Fig. 6B**). We classified these into two pair categories: Category 1 (Cat 1), representing amino acid pairs enriched in windows covering the AiSu response class; and Category 2 (Cat 2), representing pairs enriched in windows covering the AiSr class. These categories were then used to label phosphopeptides in the entire dataset irrespective of response class. Sequence logos generated from the matching Cat 1 and Cat 2 peptides revealed distinctive motif signatures that were refined from previously assessed pLogos carried out based on response class (**Fig. 6C,E**). Notably, Cat 1 phosphopeptides were enriched in -3/-2 D but also S; and +1 NLIV, suggesting phosphorylatable serine residues, in addition to -3/-2 D, can be found upstream of phosphosites whose acid-induced phosphorylation are unaffected by sorbitol. Cat 2 phosphopeptides were hyper-enriched in -3/-2 R. Strikingly, phosphopeptides harboring Cat 1 or Cat 2 motifs are distinctly distributed in the 2-dimensional MS response cloud. Where Category 1 phosphopeptides exhibited a pronounced acid response that is centered on the AiSu response class, Category 2 phosphopeptides showed a strong rightward shift indicative of canonical acid stress signaling that is repressed by sorbitol (**Fig. 6D,F**).

**Fig. 6.**
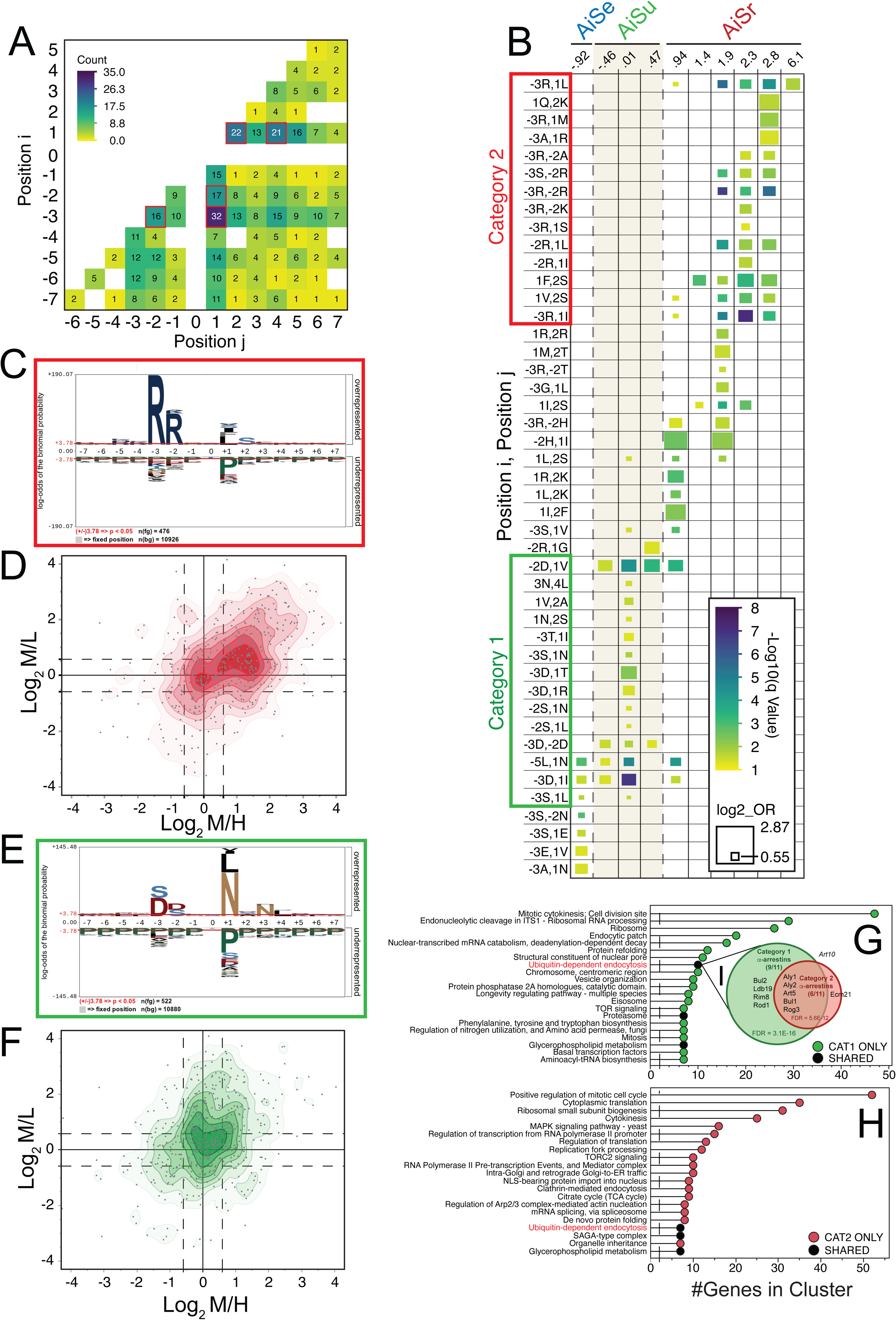
Pairwise amino acid enrichment highlights a minimal motif for AiSu and AiSr response classes. (A) Coincidence frequency of positional pairs identified as having significant amino acid pair enrichment for Log_2_M/L>0.585. Red boxed positional pairs were deconvoluted further in panels B-H. (B) Sliding window enrichment statistics for positionally-enriched amino acid pairs. Data are ploted with respect to the center of the enrichment window with the respective response classes indicated (top). Category 1 and Category 2 boundaries were defined by contiguous, multi-window enrichment spanning the AiSu (Cat 1) versus AiSr (Cat 2) response classes. (C) All phosphopeptides harboring Cat 2 amino acid pairs in B were evaluated for enrichment relative to the entire phosphoproteomic dataset using pLogos. Significant enrichment is indicated by letters reaching above the red line (Log Odds = 3.78). (D) Contour/dot plot showing the acid verus acid/sorbitol response for all phosphopeptides harboring Cat 2 amino acid pairs. (E) Same as C but for Cat 1 phosphopeptides. (F) Same as D but for Cat 1 phosphopeptides. (G) MCL-clustered gene ontologies for Cat 1 phosphopeptides. (H) MCL-clustered gene ontologies for Cat 2 phosphopeptides. (G-inset) Comparison of yeast alpha arrestin proteins harboring Cat 1 versus Cat 2 phosphosite motifs.

Gene ontology enrichment shows that Cat1 substrates are strongly associated with the bud neck/cell division machinery and polarized growth program, whereas Cat2 substrates are comparatively more associated with canonical kinase/signaling functions. For example, Cat1 substrates are enriched cytokinesis, septin ring organization, exocytosis/endocytosis, and cell polarity, which fit a program coordinating morphogenesis and cell-surface remodeling during division (**Supplemental Fig. S6A**). GO cellular component terms reinforce this clearly, with enrichment at the plasma membrane, cell division site, bud neck, septin ring, cell cortex, cytoskeleton, and vesicle-related compartments, which immediately points to proteins involved in budding, cytokinesis, membrane trafficking, and spatial organization of growth (**Supplemental Fig. S6B**). Molecular functions enriched by Cat1 substrates are associated with GTPase regulator/nucleotide-binding and enzyme regulator activities and consistent with control by small GTPases and trafficking/polarity networks rather than direct kinase catalytic output (**Supplemental Fig. S6C**). By contrast, the terms biased toward Cat2 are fewer and generally broader, including kinase activity, phosphatidylinositol/phospholipid binding, and broader cell-cycle regulatory terms consistent with Cat2 representing canonical acid-induced TORC2/PKC-linked signaling that has broad impacts across the cell.

Gene cluster analysis further showed that Cat1 and Cat2 share a core growth-control network involving membrane trafficking, lipid homeostasis, proteostasis, and regulatory complexes, but they differ in emphasis. Cat1 substrates are more strongly associated with budding, cytokinesis, polarized growth, and the cortical/endomembrane machinery that drives cell morphogenesis, whereas Cat2 substrates are more abundant in signaling, translation, transcriptional control, stress-response pathways, and broader homeostatic regulation (**Fig. 6G&H**). In general, Cat1 substrates appear linked more to the execution of growth and division at the cell cortex and bud neck, while Cat2 substrates are linked more to the regulatory systems that coordinate those processes. Notably, Cat1 substrates were also more prominently enriched than Cat 2 in ubiquitin-dependent endocytosis, including 9 out of the 11 yeast alpha arrestins, which serve as E3 ligase adaptors for the endocytic trafficking of plasma membrane proteins like the G protein coupled receptor Ste2 (28) (**Fig. 6I**). No other gene cluster showed as much coverage of a GO term, suggesting a unique connection between the Cat1 motif and alpha-arrestins.

Taken together, these results reveal a minimal motif and distinctive mechanistic roles for non-canonical acid-dependent TORC2/PKC-independent kinase signaling.

### Casein kinase, Yck1, is a primary driver of non-canonical acid stress phosphorylation

Independently from our analysis of sequence motifs, we looked for enrichment of known kinase/substrate pairs across all response classes. To do this, we used the BioGRID database to identify known kinase/substrate pairs in our dataset and then asked whether any pairs were significantly enriched in one of the six response classes. Pairs were defined at either the *site level*, wherein a kinase/pSite relationship is known, or at the *substrate level*, wherein a known kinase/substrate relationship is tested for enrichment regardless of pSite pairing. Response-class enrichment at the site level did not yield any technically significant results (-Log_10_ q < 1.3), due largely to a lack of data in general (**Fig. 7A, Supplemental Fig. S7**). In comparison, significant enrichment was observed at the substrate level for multiple response classes. In particular, Psk2, Yck1, Ypk1 and Pkh3 were enriched in the AiSu response class compared with Rck2, Kkq8, Ssn8, Qrk1, Pkh1, and Npr1 in the AiSr class (**Fig. 7A**). Sliding window analysis of kinase/substrate pairs highlighted distinct enrichment trends for each kinase that were reminiscent of the trends in motif enrichment (**Fig. 7B,C**; compare with Fig. 4G-I). However, in some cases the kinase enrichment was inconsistent with motif enrichment, likely because substrate level enrichment analysis does not discriminate between specific pSites. For example, Pkh3, Hrk1, and Ypk1, which are enriched in AiSu, are basophilic rather than acidophilic kinases. Thus, these cases may represent crosstalk at the substrate level that convolutes multiple kinases in the analysis. Most other kinases were either basophilic and consistent with enrichment in AiSr, +1P sites that follow CDK logic, or kinases with poorly defined consensus sequences such as Pkh3.

**Fig. 7.**
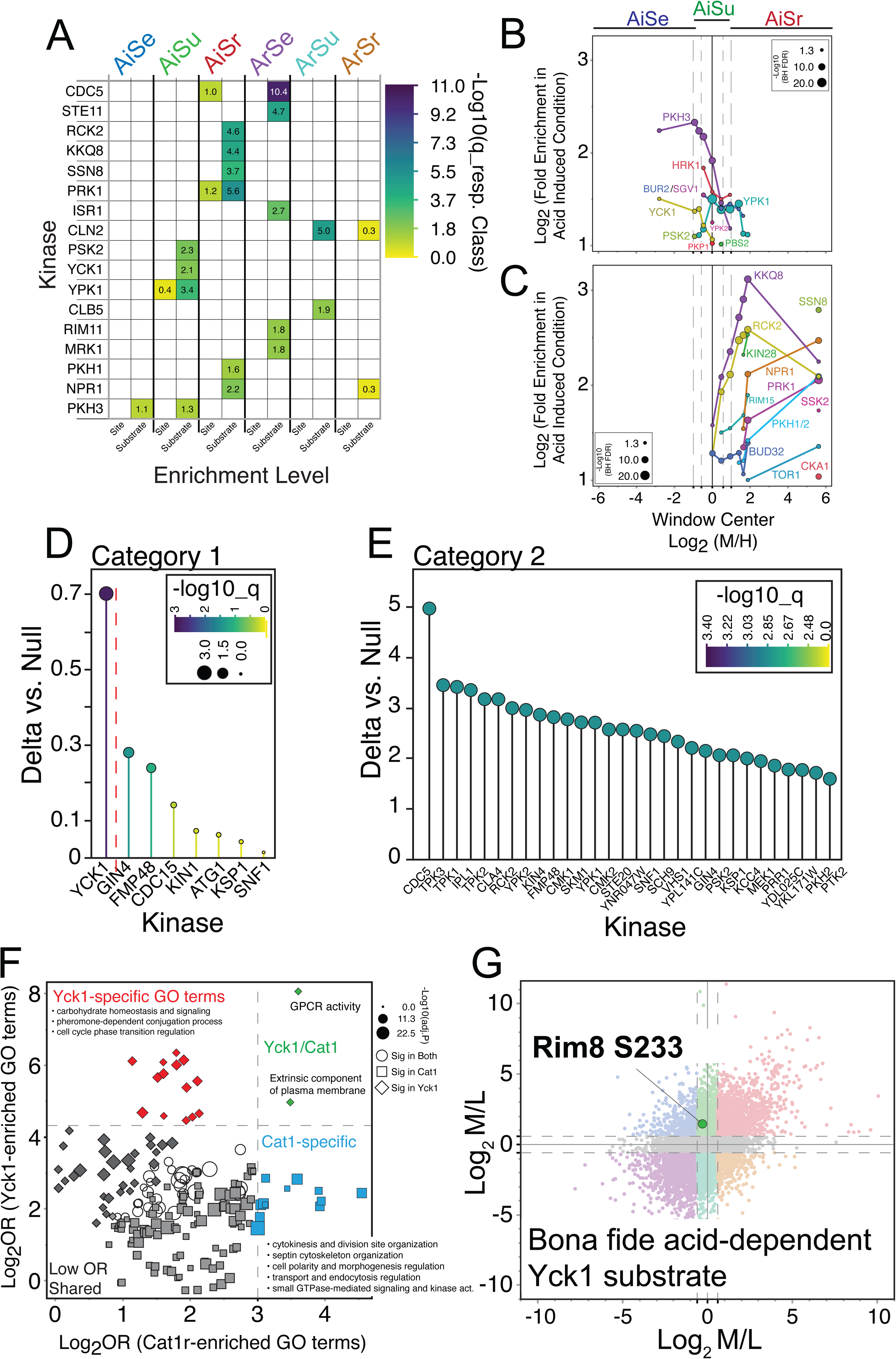
Kinase enrichment highlights Yck1 as the likely driver of a TORC2/CWI-independent acid stress response. (A) Kinase-to-phosphosite (site) or kinase-to-protein substrate (substrate) pairs were identified via BioGRID and tested for enrichment in the acid response classes. Shown are the log_10_(q-Value) for enrichment in the region relative to the entire phosphoproteome dataset. Values <1.3 represent q<0.05. (B,C) Windowed enrichment trends for kinase substrates in the acid induced half of the response. To improve graph clarity, the plots are separated based on trend and overlap with AiSu versus AiSr response classes. (D) Kinase-to-substrate motif matching using probability weight matrices (PWMs). Match quality is indicated by the delta vs null score representing the response category 1-specific kinase motif preference relative to the global phosphosite background while significance of the match was achieved via permutation testing. (E) Same as in D but for Cat 2 sequences. (F) Shared versus distinctive gene ontology enrichment for Cat 1 versus Cat 2 sequences as determined using g:Profiler multi-comparisons and log odds normalization. Color coding indicates groupings arbitrarily defined by natural Yck1-specific, Cat 1-specific, and shared GO terms. (G) Rim8 alpha arrestin phosphorylation site S233, shown within the context of our phosphoproteomics dataset, was observed in our dataset and is established to be co-dependent Yck1 and acid stress.

To deconvolute the kinase/motif relationships in our dataset we adapted previously published position weight matrices (PWM) that quantify the specificity of yeast kinases for any given substrate sequence based on in vitro peptide array data (29) (see Methods). Cat 1 and Cat 2 phosphopeptide sequences were then scored in the context of the average PWM score per kinase relative to the average score across all kinases (represented as a delta vs. null score) followed by permutation testing. Results highlighted prominent enrichment of the acidophilic membrane-anchored kinase, Yck1, as the only significant match for Cat 1 sequences (**Fig. 7D**). In comparison, several kinases were significantly enriched for the Cat 2 sequences including AGC kinases, YPK1/2, and several other kinases whose substrates are enriched in the AiSr class (**Fig. 7E** and **7C**).

We next asked whether the known Yck1 protein network overlapped with protein networks defined by Cat 1 sequences with acid induced phosphorylations. Canonical Yck1 protein networks are enriched in carbon source homeostasis and signaling gene ontologies reflective of its well-published roles glucose sensing (**Fig. 7F**, red diamonds). Strikingly, Cat 1 sequences show enrichment of several GO terms that were also enriched by Yck1 itself. Beyond the generic terms, we noticed significant enrichment of MAPK signaling, plasma membrane, cellular bud neck and site of polarized growth (**Fig. 7F**, open circles; **Supplemental Fig. S8A**). We also found high odds ratio enrichment of the terms GPCR activity and extrinsic component of plasma membrane for both Yck1 and Cat1 proteins, in part reflecting the published evidence of Yck1’s role in ubiquitin-mediated endocytosis of pheromone receptor Ste2 (**Fig. 7F**, green diamonds) (30). Cat 1 sequences also showed high odds ratio enrichment of GO terms that exhibit relatively lower odds ratios for Yck1 protein networks that are annotated currently. These included proteins involved in cytokinesis, cytoskeletal organization, cell polarity, and endocytosis (**Fig. 7F**, blue squares; **Supplemental Fig. S8B**). Not surprisingly, many of Cat 1-associated terms that were shared with Yck1 protein networks were also observed in Cat 2-associated protein networks but were not specifically enriched (**Supplemental Fig. S8C**). In conclusion, acid-induced Cat1 phosphosite-containing protein networks overlap significantly with Yck1-specific protein networks that support their potential service as Yck1 phosphosites.

We next sought to find evidence for acid-activated Yck1-dependent phosphorylation that could confirm our findings. Annotation of kinase/pSite pairings remains a significant challenge since the substrate preferences of kinases are rarely discrete and publications rarely contain kinase/substrate studies that are acutely resolved at the amino acid site level (31). Not surprisingly, BioGRID contained only 3 Yck1 kinase/substrate pairs that overlapped with our dataset, only one of which presented useful biochemical evidence, although the evidence was not resolved at the site level. However, text mining of the published literature produced two notable cases, both of which highlighted several consistencies with our data. Case one involves Yck1-dependent phosphorylation of the proton pump Pma1, which occurs at the plasma membrane, favors -3D phosphosite sequences, and was shown to increase in response to lowering the pH regardless of the presence or absence of sorbitol (32). In case two, Yck1-dependent phosphorylation of the α-arrestin Rim8 at the plasma membrane occurs on an acidophilic phosphosite in an acid-titratable manner (33). Of these two published examples, only Rim8 phosphorylation was mapped to a precise site previously (Ser233), which we also observed as acid-induced/sorbitol unaffected (AiSu) in our dataset (**Fig. 7G**). Other substrate proteins have been identified for Yck1 but their sites of phosphorylation have not been investigated under acid stress conditions, nor have they been sufficiently distinguished from substrates of the paralog, Yck2, which is commonly nullified in tandem. These results provide supporting evidence for Yck1 as one of the key drivers of acid-induced, TORC2/PKC-independent phosphorylation in yeast.

## DISCUSSION

We identified subsets of the yeast phosphoproteome whose responses to acetic acid stress are either dependent on or independent of canonical acid stress signaling through the TORC2 and CWI pathways, which are widely thought to be coupled to changes in membrane structure. TORC2/CWI-independent, acid-induced phosphorylation occurs on a distinct network of proteins with cellular localizations, biological processes, and molecular functions that differ from those associated with TORC2/CWI-dependent signaling. Notably, many of these sites are acidophilic phosphosites enriched at the plasma membrane, consistent with the sequence and localization preferences of substrates of Yck1, a membrane-anchored casein kinase previously shown to phosphorylate acidophilic sites in response to acetic acid. In contrast, TORC2/CWI-dependent substrates are predominantly basophilic and linked to several kinases involved in TORC2 and CWI signaling. We also found additional local sequence constraints of acid stress regulated phosphosites. Most prominently, phosphorylation of proline-rich sequences is quantitatively repressed in acid stress while phosphorylation of histidine-rich sequences is enhanced. Finally, our findings suggest that TORC2/CWI-independent, acid-induced phosphorylation is strongly associated with G protein signaling and ubiquitin-mediated endocytosis, processes with well-established connections to yeast budding, cytokinesis, cell polarization, yeast mating projections and Yck1 (34–37). All together, these findings support the existence of a previously underappreciated arm of pH stress signaling that is driven in part by Yck1 and distinct from canonical TORC2/CWI pathway output (**Fig. 8**).

**Fig. 8.**
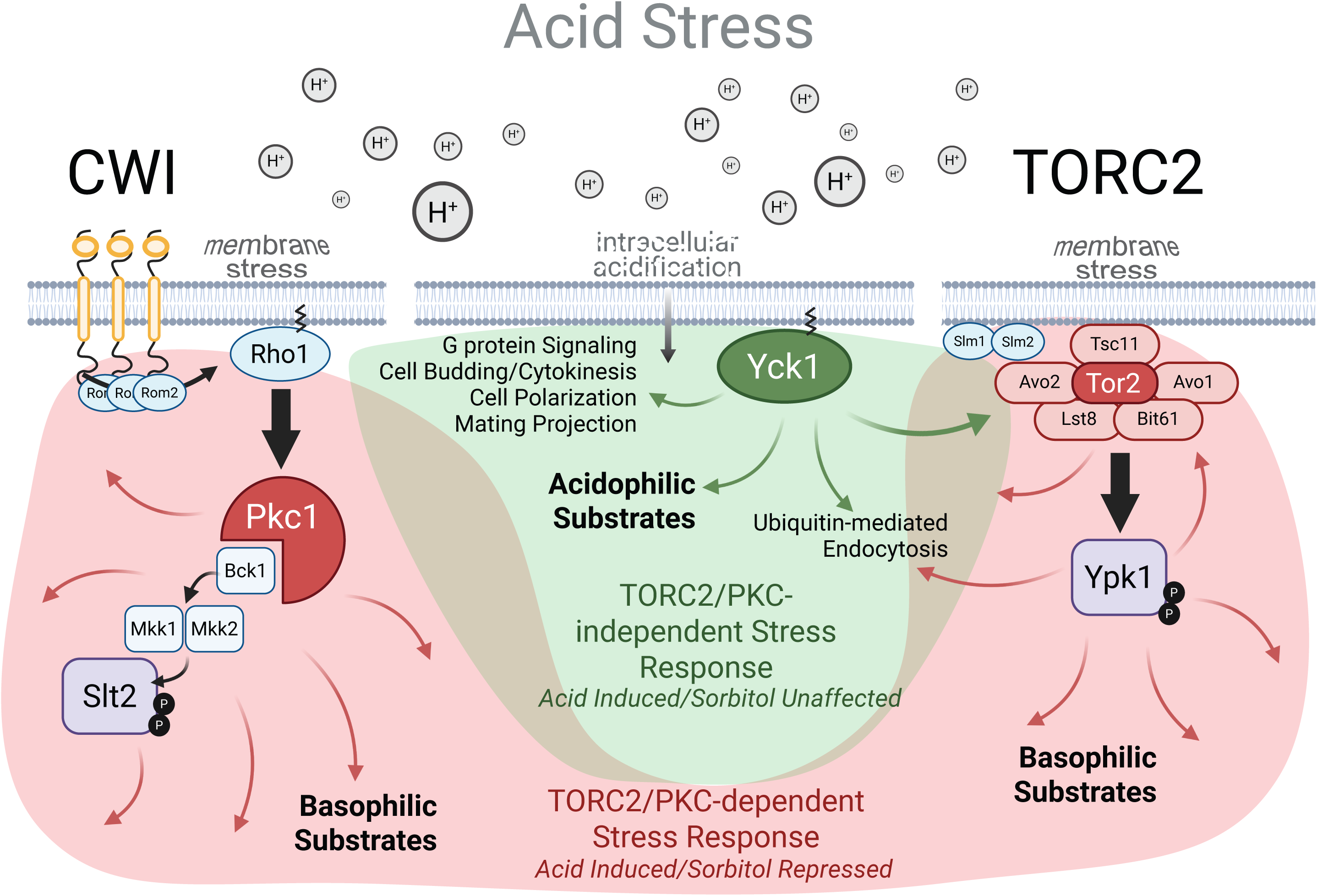
Acetic acid stress activates three major signaling responses. Acetic acid stress elicits a broad range of changes in the phosphoproteome of yeast. Canonical pathways of the acetic acid stress response include the cell wall integrity and TORC2 pathways that are activated by changes in acid-dependent membrane stress and mediated through Pkc1 and Tor2/Ypk1 kinases, that promote phosphorylation of basophilic kinase substrates (often -3/-2R) (red shading). Activation of these two pathways propagates to other kinases and substrates (curved arrows). A non-canonical, third pathway is activated in tandem, which is independent of membrane stress and likely due to intracellular acidification (green shading). This pathway is marked by the phosphorylation of acidophilic kinase substrates localized at the plasma membrane, such as the GPCRs, G proteins, proteins involved in cell polarization, and cell budding. This non-canonical pathway is independent of membrane stress pathways and is likely mediated through activation of the membrane-localized casein kinase, Yck1. Considerable overlap occurs between protein substrates of both the canonical and non-canonical paths leading to an integrated response to the acid stress.

### Comparison with previous phosphoproteomic studies of acid stress

Our study is the first to dissect acidic pH stress effects on the phosphoproteome. However, it is not the first to carry out a phosphoproteomic analysis of acetic acid or otherwise low pH stress condition. Although not shown here, we carried out exhaustive searches for any and all pH-associated kinases to explain our data. Notably, Guerreiro et al. carried out an LCMS analysis of the yeast membrane phosphoproteome, which highlighted Hrk1 kinase as a predominant driver of acetic acid-dependent phosphorylation of membrane-localized lipid biosynthetic pathways (14). We utilized phosphosites identified from that study to track down Hrk1 substrates in our dataset (which were not available on BioGRID), finding that they too are enriched in the AiSu response class (**Fig. 7B**). However, there is no well-defined consensus motif for Hrk1 substrates and we were unable to extract a reliable motif that could be used to explain the bulk of our data. Nonetheless, Hrk1 is also a known kinase for Pma1 among other plasma membrane transporters, suggesting its involvement with Yck1 in the AiSu response.

A large-scale analysis of yeast conditional phosphoproteome responses by Leutert et al also captured the effects of low pH stress (38). Somewhat similar to our study, they found an LN(K/R)NNpS(F/M)xDL motif with clear enrichment of -5L, +4L, and asparagine rich sequences. However, they observed only basophilic sequence enrichment that is likely to have overwhelmed any acidophilic substrates in the absence of a sorbitol blockade like the one we performed. Indeed, we observed a similar pLogos enrichment for the Ai response class when ignoring the sorbitol effects. The consistencies between the two datasets suggest some importance of leucine in proximal positions that are not yet well understood and that we have not investigated here.

### Acidophilic substrates: A core network motif in the TORC2/PKC-independent acid response?

While we were able to identify acidophilic substrates with -3/-2D enrichment, they represent only a fraction of the total phosphosites observed in the AiSu class. We made every attempt to extract yet additional motifs from the data, including several different parsing strategies that separate the data beyond 2 dimensions. However, none were able to identify any other statistically enriched motifs. Beyond using pLogos, which only captures single position enrichments, we established pairwise relationships in neighboring amino acid positions that further restrict the set of acid induced and sorbitol unaffected phosphorylation sites we defined as category 1 sites (**Fig. 6**). Projecting all category 1 phosphopeptides back onto the phosphoproteome response grid revealed the best of all observed outcomes for any AiSu motif, showing very good agreement with the AiSu response class. Due to the variability in neighboring residues, we were unable to achieve this same degree of motif-to-response agreement by any other means, including several published clustering, motif, or MMD permutation algorithms such as CDhit, SELPHI2, or Gibbs Cluster (39–41). We also attempted a subtraction method for identifying other potentially hidden motifs to no avail. We hypothesize that the acidophilic motif was observable because it represents a core acid response network for the AiSu response class, which becomes buried in the noise of promiscuous tertiary and quaternary phosphorylations after 15 minutes of acid treatment. Future work might deconvolute this further via temporal analysis, which was shown previously to delineate between early-stage primary/core response networks and late stage promiscuous phosphorylation networks in the high osmolarity/glycerol pathway (42).

### Hyperphosphorylation of histidine-rich sequences: Support for the hypothesis that protons act as second messengers to conditionally activate kinase networks

We made a robust observation that phosphosites harboring histidine at positions -4 and -3 are hyperphosphorylated on average relative to any other amino acid in the first five positions preceding the phosphosite (**Fig. 5**). This pattern was detected by exclusive analysis of the extreme outlier phosphosites and was only found in the AiSr respsonse class, which harbors basophilic kinase substrates (-3/-2 R). We submit that this supports the hypothesis that protons act as second messengers to conditionally activate kinase substrate networks. This hypothesis is further supported by a previous in vitro study demonstrating that a -3H phosphosite-containing peptide in the pyrimidine biosynthesis multienzyme CAD becomes a preferred and rapidly catalyzed substrate for cyclic-AMP PKA under low pH conditions (43). Comparatively, phosphorylation of a canonical -3R phosphosite-containing peptide was unaffected by titrating pH conditions. Further analysis showed that these effects were independent of structural changes and therefore more likely due to histidine protonation. These data support the hypothesis that protonated histidine at the -4 or -3 position relative to a phosphosite may activate some substrates for phosphorylation by basophilic kinases. Considering that exposed histidine residues exhibit a pKa ∼8, whereas the pKa of a buried histidine is between 5.5-6, may further pinpoint His-containing phosphosites in highly exposed disordered regions as potential targets for regulation by hyperphosphorylation under acid stress.

Beyond the substrate, prior evidence has shown that protonation of histidine can also serve as a pH-governor of enzyme activity (11, 44–46). This has recently been shown for polyHis tracts that regulate Snf1, a kinase that is sensitive to glucose starvation, which also lowers intracellular pH (47).

### Poly-proline is generally antagonistic to acid-stress phosphorylation

A striking trend in our dataset was the quantitative anti-correlation between acid stress phosphorylation and localized proline content (**Fig. 4** & **Supplemental Fig. S4**). This trend extended between 1 to 7 proline residues and therefore cannot be explained by a regulatory inhibition mechanism of +1P substrates that are targets of cyclin dependent and mitogen activated protein kinases. Indeed, several +1P substrates exhibit elevated phosphorylation in response to pH stress (**Supplemental Fig. S4**). We are unaware of such an observation being made previously. Unlike histidine, the sidechain of proline is not ionizable under normal biological conditions and therefore protonation is unlikely a factor. Alternatively, cis/trans proline isomerization, which can be influenced by pH, may provide a more plausible cause for the trend due to changes in local protein structure that could potentially change kinase accessibility (48–51). Nonetheless, our observation can contribute to sequence-based algorithms designed to predict phosphorylation behavior.

### Yck1, casein kinases, acidophilic substrates, and TORC2/CWI-independent acid stress signaling

Our data support the hypothesis that Yck1 is a primary driver of the acid-dependent, sorbitol-insensitive phospho-response to acetic acid stress. This conclusion is supported by condition-dependent substrate motif enrichment, PWM-based motif scoring, substrate localization, co-enrichment of known substrates, gene ontology co-enrichment, and prior evidence linking kinase activity to acid stress. To our knowledge, this is the first proteome-wide evidence identifying Yck1 as a central kinase in the acetic acid response, made possible by separating this pathway from membrane integrity-dependent TORC2 and CWI signaling. Consistent with this distinction, canonical TORC2/CWI substrates are basophilic, whereas the Yck1-associated substrates we observe are strongly acidophilic.

Yck1 is membrane anchored, placing it near established acid-dependent substrates such as Pma1 and Rim8, as well as glucose-regulated substrates including Rgt2, Snf3, Mgt1, and Std1, all of which localize to the plasma membrane (32, 33, 52, 53). However, the mechanism by which Yck1 is activated remains unresolved. Although Yck1 has established roles in both glucose- and pH-responsive signaling, these may reflect distinct modes of regulation. Glucose starvation causes only a modest decrease in intracellular pH, whereas the 45 mM acetic acid condition produces more pronounced intracellular acidification (10, 15, 54). Consequently, the mechanisms of Yck1 activation in these two scenarios may not be the same. A study looking at the Yck1-phosphorylated C-terminus suggests a more constitutive, rather than dynamically regulated, behavior of the kinase is involved in control of glucose transporters Rgt2 and Snf3 (55, 56). Moreover, it’s been shown that glucose receptors are epistatic to Yck1, arguing against the premise that the kinase is activated by glucose receptors (52).

A major obstacle in defining Yck1 signaling is the lack of site-resolved phosphorylation evidence. In prior studies of glucose-dependent Yck1 activity, target phosphosites were often inferred rather than mapped directly. When we examined these reported sites, many lacked the upstream acidic residues characteristic of casein kinase substrates, and none were detected in our dataset. We did, however, observe several alternative phosphosites in Yck1 substrates that were differentially phosphorylated in acid stress (see supplemental tables). These included a mixture of local sequence that were insufficient to highlight any one kinase. Our evidence reveals hundreds of acidophilic phosphosites induced by acid stress that are unaffected by sorbitol, and conform to bona fide casein kinase phosphosites including those shown previously to be direct acid dependent substrates of Yck1 (32, 33). We suggest that these represent a TORC2/CWI-independent signaling network that has not yet been mapped in the gene ontology databases, which would explain why Yck1 associated networks only partially overlap with acidophilic Cat 1 phosphosite-containing protein networks (**Fig. 7F** & **Supplemental Fig. S8**).

### How is Yck1 activated to phosphorylate its substrates?

The activating mechanism for Yck1 remains elusive. What has been studied is usually in the context of glucose signaling. Early studies presented evidence that the glucose sensors are coupled to membrane-associated Yck1 (57). However, that model was weakened by later work showing that direct interaction between the glucose sensors and Yck1/2 is not required for glucose-induced degradation of the glucose sensor co-repressor Mth1, and suggested that the signal may instead pass through a yet unidentified component (58). There is some evidence for upstream control of Yck1/2 by phosphoregulation. Gadura et al. concluded that Glc7-Reg1 phosphatase acts as an upstream activator of Yck1/2 in carbon-source signaling, consistent with Yck1 phosphorylation state being regulated (59). We observed several Yck1 phosphosites in our dataset, most of which were acid induced and either unaffected or only mildly repressed by sorbitol, consistent with the hypothesis that they could be activating under low pH and independent of TORC2/CWI pathways (see supplemental tables). However, the local sequences of these are mixed and do not point clearly to any one kinase. Beyond extrinsic phosphorylation, other casein kinases are known to self-regulate through auto phosphorylation, which may also play a role in conditional Yck1 activity control. Thus, the mechanisms of Yck1 activation under acetic acid and glucose stress remain an outstanding question. This work can help by providing a foundation for comparison to Yck1-specific studies in the future.

### Limitations of the study

In this study, we used a cell wall stabilizer, sorbitol, to selectively block the activation of TORC2 and CWI pathways, thereby allowing us to focus on the intracellular acidification effects of the stress response. Experimental validation confirmed the blockade of cell wall-related signaling pathways such as cell wall integrity pathway and TORC2-Ypk1 signaling. Sorbitol, known to increase extracellular osmolarity, induces the activation of the high-osmolarity glycerol (HOG) pathway in yeast, but the activation will not persist beyond 1.5 hours as reported in the literature and confirmed in our experiments (**Fig. S9**, without sorbitol condition) (60–62). This is an important observation, indicating that our equilibration of the yeast in sorbitol before acid treatment should not lead to confounding stress responses that would confound the dissection of the effects of intracellular acidification. In summary, sorbitol allows the blockage of cell wall stress-related signaling without inducing other stress responses or metabolizing of the compound (63).

One limitation of approach is the potential for acetic acid to induce feedback responses after blocking cell wall-related pathways. For example, we observed an acid-induced activation of Hog1 only in the sorbitol supplement condition, indicating feedback effects of suppressing cell wall stress-related signaling on the HOG pathway (**Fig. S8**). This crosstalk has been observed before in response to zymolyase (64, 65). Considering the crosstalk between CWI and other signaling pathways, other feedback effects may also exist (66–68). Fortunately, this effect is not expected to significantly interfere with the identification of pH_i_-sensitive phosphorylations as they are unaffected by sorbitol treatment. Other categories, such as sorbitol-enhanced (Se) phosphorylation may contain phosphorylations directly induced by sorbitol and/or influenced by the feedback of CWI inhibition. Under this scenario, validation of phosphorylation level by pH_i_ titration may further confirm the pH_i_-sensitive phosphorylations identified in this study are solely and directly responsive to intracellular acidification.

## METHODS

### Yeast cell culture and SILAC labeling

SC-Lys-Arg media supplemented with an appropriate amount of heavy medium or light lysine and arginine was used for cell culture. 1.7g/L yeast nitrogen base (YNB) without amino acids without ammonium sulfate, 5g/L ammonium sulfate, and 2.0% (w/v) glucose, supplemented with selected amino acids and nucleobases (L-histidine, L-tryptophan, L-methionine, adenine, and uracil, 20 mg/L each; L-isoleucine and L-tyrosine, 30 mg/l each; L-phenylalanine, 50 mg/l; L-leucine, 100 mg/l; L-valine, 150 mg/l; L-threonine, 200 mg/l; L-proline, 200 mg/L; L-lysine and L-arginine, 50 mg/l each); ‘light’ medium is supplemented with regular Lys and Arg; ‘medium’ medium is supplemented with Lys4 and Arg6; ‘heavy’ medium is supplemented with Lys8 and Arg10. lys1*Δ* arg4*Δ* (MATa his3*Δ*1 leu2*Δ*0 met15*Δ*0 ura3*Δ*0 lys1*Δ* arg4*Δ*) single colonies were inoculated in 3mL synthetic complete (SC) start culture and grown to saturation. Next day, cultures were diluted to OD600 0.1, grown until OD600 ∼0.8 in SC media, and then seeded at OD 0.00075 in SC media containing heavy, medium or light lysine and arginine, respectively. SC heavy media was supplied with 1M sorbitol. Cells were grown to the mid-log phase, and then heavy and medium cultures were treated with 45 mM acetic acid for 15 minutes. After treatment, cell growth was arrested by the addition of TCA to 2.5%. After this, cells were washed with chilled water, pelleted, and stored at -80C. Two biological replicates were analyzed at each condition.

### Cell lysis, protein extraction and digestion

Cells were lysed in 6 M urea, 75 mM NaCl, 50 mM Tris-HCl buffer (pH 8.2) using glass beads by vertexing at 4°C for 10 minutes. Lysates were centrifuged and collected supernatant. Protein concentrations were determined by DC protein assay, and heavy, medium, and light samples were mixed at 1:1:1 ratio. Mixed samples were reduced with 5mM TCEP in the dark at room temperature for 1 hour and alkylated with 10 mM iodoacetamide (IAA) in the dark at room temperature for 30 minutes. Trypsin was added at a final protease: protein ratio of 1:50 (w/w) and incubated with samples at 37°C for 18 hours. Digested samples were desalted through Sep-Pak tC18 cartridges and lyophilized by CentriVap.

### High-pH RPLC pre-fractionation

Peptide mixtures were fractionated using Waters XBridge C18 RP column (4.6 x 250mm, 3.5 µM) on Agilent 1100 HPLC operating at 0.7 mL/min as described previously. Buffer A consisted of 10 mM ammonium formate and buffer B consisted of 10 mM ammonium formate with 90% acetonitrile were used at pH 10. The fractionation gradient commenced as follows: 5% B to 25% B in 50 min, ramped to 75% B in 10 min and held at 75% B for 5 minutes. At this point, fraction collection was halted, and the gradient was ramped back to 5% B, where the column was then washed and equilibrated. Collected samples were further combined into 13 fractions for phosphopeptides enrichment and 21 fractions for proteome analysis.

### Phosphopeptide enrichment

Phosphopeptides were enriched using High Select™ Phosphopeptide Enrichment Kits (Thermo Scientific™) and followed the manufactory protocol in each of 13 fractions.

### LC-MS/MS analysis

Lyophilized phosphopeptides were resuspended in 0.1% formic acid in 10% acetonitrile and then analyzed independently by LC-MS on two separate mass spectrometers. The first analysis was carried out using a Q-Exactive Plus orbitrap mass spectrometer equipped with Dionex UltiMate 3000 LC system (Thermo). Briefly, peptides were loaded onto a trap column (PepMap™ NEO 5 µm C18 300 µm X 5 mm Trap Cartridge) and resolved through a custom analytical column packed with ReproSil-Pur 120 C18-AQ 3 µm beads (Dr. Maisch GmbH) at a flow rate of 0.3 µL/min with a gradient solvent A (0.1% formic acid in 2% acetonitrile) and a gradient solvent B (0.1% formic acid in 80% acetonitrile) for 150 minutes. MS analysis was conducted in a data-dependent manner with full scans in the range from 400 to 1800 m/z using an Orbitrap mass analyzer set as follows: MS1: resolution = 70,000, AGC target = 3e^6^, Max IT = 100ms; MS2: resolution = 17,500, AGC target 1e^5^, Max IT = 50ms. The top fifteen most intense precursor ions were selected for MS2 with an isolation window of 4 m/z. Isolated precursors were fragmented by high energy collisional dissociation (HCD) with normalized collision energy (NCE) of 27.

The second LCMS analysis was carried out using a timsTOF HT mass spectrometer coupled with a nanoElute 2 UPLC system equipped with c18 analytical UPLC column (Ion Opticks 25cm x 75μm, 1.7μm) held at 50C using a Bruker column Toaster (Bruker Daltonics Inc.). Peptides were separated using a 30 minute gradient of buffers A (0.1% formic acid in water) and B (0.1% formic acid in acetonitrile): 5%-26% B from 0-14 minutes at 0.25 μl/min; 26%-40% B from 14-22 minutes at 0.25 μl/min; 40%-90% B from 22-26 minutes at 0.35 μl/min. The TIMS module was set to custom dda-PASEF mode with the following parameters: 1/K0 start = 0.64; 1/K0 end = 1.4; Ramp & Accumulation time = 50 ms; Duty cycle = 100%; ramp rate = 17.53 Hz. Peptides were analyzed from a mass range of 349 Da to 1234 Da and fragmented using scaled collision energy from 20 eV at start to 59 eV at end.

### MS data processing

Orbitrap RAW files were searched against the yeast proteome sequence database from UniProt using the Sequest HT search engine embedded in Proteome Discoverer 3.0 (Thermo) with 10 ppm MS1 precursor mass tolerance, 0.1 Da MS2 fragment mass tolerance, 0.01 false discovery rate. SILAC ratios were calculated based on the ratio of integrated peak areas at MS1 for SILAC ‘heavy’, ‘medium’ and ‘light’ peptides. Phosphorylation changes were normalized with protein abundance changes.

Raw timsTOF HT data acquired by DDA-PASEF and dia-PASEF were analyzed in FragPipe/MSFragger using the *S. cerevisiae* reviewed UniProt database with appended reversed decoy sequences. Searches were performed with 20 ppm precursor tolerance, 0.05 Da fragment tolerance. Fully tryptic peptides were required, with up to two missed cleavages, peptide lengths of 7–40 amino acids, and precursor masses of 500–5,000 Da. PTMProphet was used for phosphosite localization, MSBooster was used for rescoring with predicted retention time and fragment spectra, and ProteinProphet was used for protein inference. Results were filtered to 1% false discovery rate using a target-decoy approach. The statistical significance of phosphosites with acid response (medium/light) and sorbitol response (medium/heavy) was determined using response screening in JMP 16. Phosphopeptides with a single phosphorylation sites were included in the formal analyses regardless of significance thresholds unless otherwise stated.

### Gene ontology enrichment

Gene ontology enrichment analysis was carried out using STRING or g:Profiler using *Saccharomyces cerevisiae* as the background reference (69, 70). Data extracted from g:Profiler multi-queries were converted into Log_2_ of the odds ratios for figures. Data were then filtered by false discovery rates less than 5% unless otherwise shown.

### Sequence analyses

Sequence logos were generated using pLogo with 15-mer phosphosite-centered sequences (71). Residues missing at protein termini were padded with “x”. Non-redundant foreground sequences were removed from the background of 11,402 non-redundant phosphosites in our dataset before analysis. Positional physiochemical property-based enrichment was carried out using a custom Python script with the following relationships: Acidic-DE; Basic-RKH; Polar-STNQ; Hydrophobic-AVLIMFYW; Special-CPG. Windowed enrichment of individual amino acids was carried out using a custom python script designed to calculate the log odds for enrichment at any position within +/-7 of the phosphosite relative to the entire phosphoproteomic dataset.

Pairwise amino acid enrichment was carried out using a custom python script to calculate a fisher’s exact test (one-sided enrichment), log_2_ odds ratio, and Benjamini-Hochberg false discovery rate correction for windowed foreground 15mer sequences compared against the entirety of all peptide sequences detected (19,011). These were then plotted with respect to the center of the window for display purposes.

### Kinase/Substrate Analysis

Kinase/substrate pairings were downloaded from BioGRID (72) and processed using custom python scripts designed to test the enrichment of kinases within a response class or windowed response class across Log_2_(M/H) relative to the entire dataset. Enrichments were carried out at the site level, in which a precise kinase-to-pSite association was required for positive enrichment, or at the substrate level, in which kinase-to-substrate protein association was sufficient to claim positive enrichment. Significance of enrichments were estimated by fisher’s exact test (one-sided enrichment), log_2_ odds ratio, and Benjamini-Hochberg false discovery rate correction.

### Kinase/Motif Matching with Probability Weight Matrices

See extended methods supplement.

### AlphaFold Structural Comparisons

See extended methods supplement.

## ACKNOWLEDGEMENTS

We would like to thank our funding sources for support of this research, including: NSF-Simons grant 1764406 to M.T., the Georgia Tech Abell Professorship to MT, and NIH grant S10OD038327 to MT, which supports shared instrumentation access to the GT timsTOF HT. We are grateful to Griffith Carr and Diego Assis (Bruker Daltonics) for support with timsTOF HT mass spectrometry analysis. We would also like thank Daniel Isom (U. Miami) and Josh Kelley (U. Maine) for critical review and comments.

## SUPPLEMENTAL INFORMATION

**Supplemental Fig. S1.**
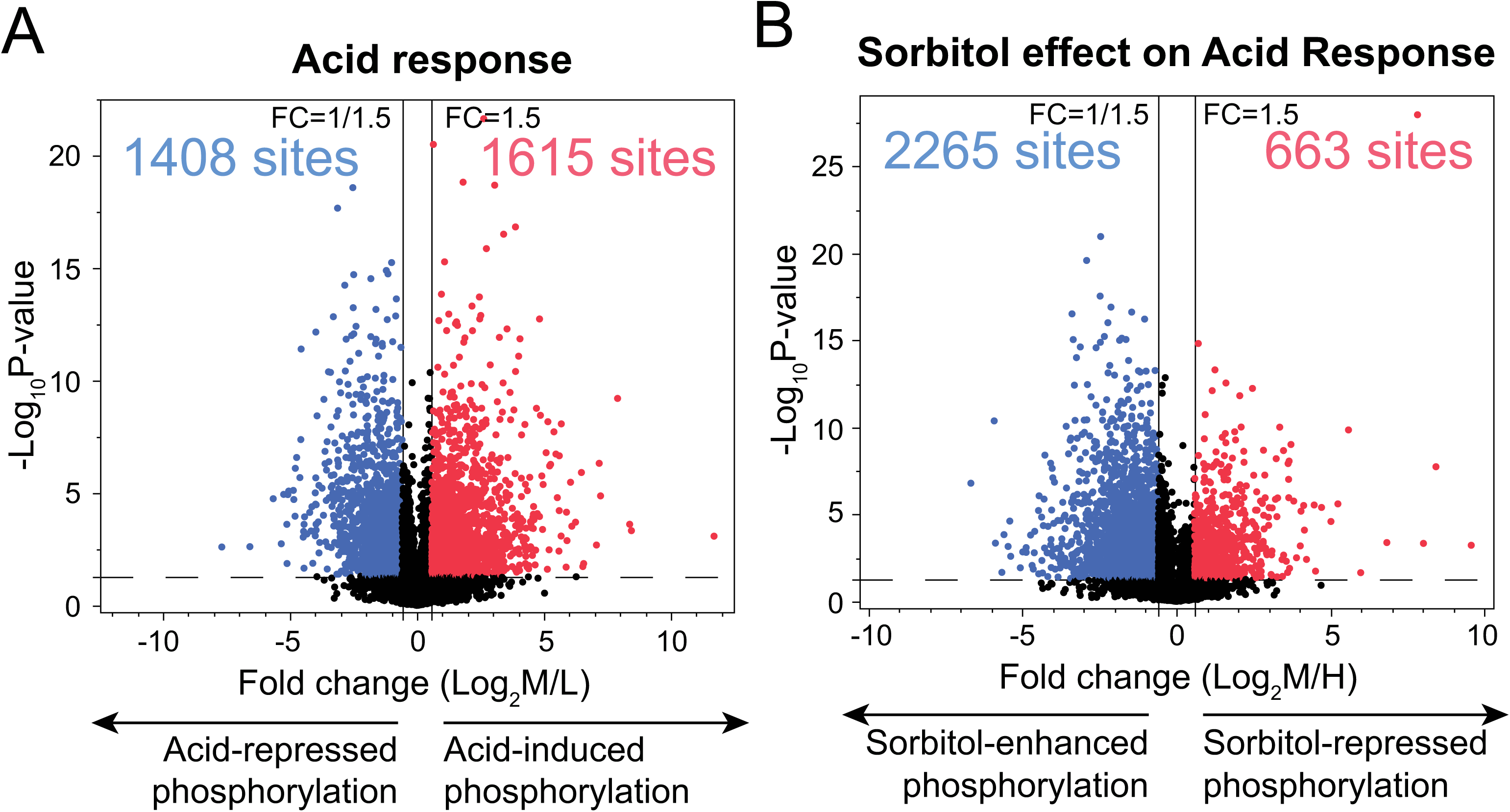
(A) Volcano plot for acid vs control (Log_2_M/L). (B) Volcano plot for sorbitol-equilibrated acid vs control (Log_2_M/H). (Dashed horizontal line) Significance threshold p = 0.05. (Solid vertical lines) Log_2_ responses representing a -1.5 and 1.5-fold change.

**Supplemental Fig. S2.**
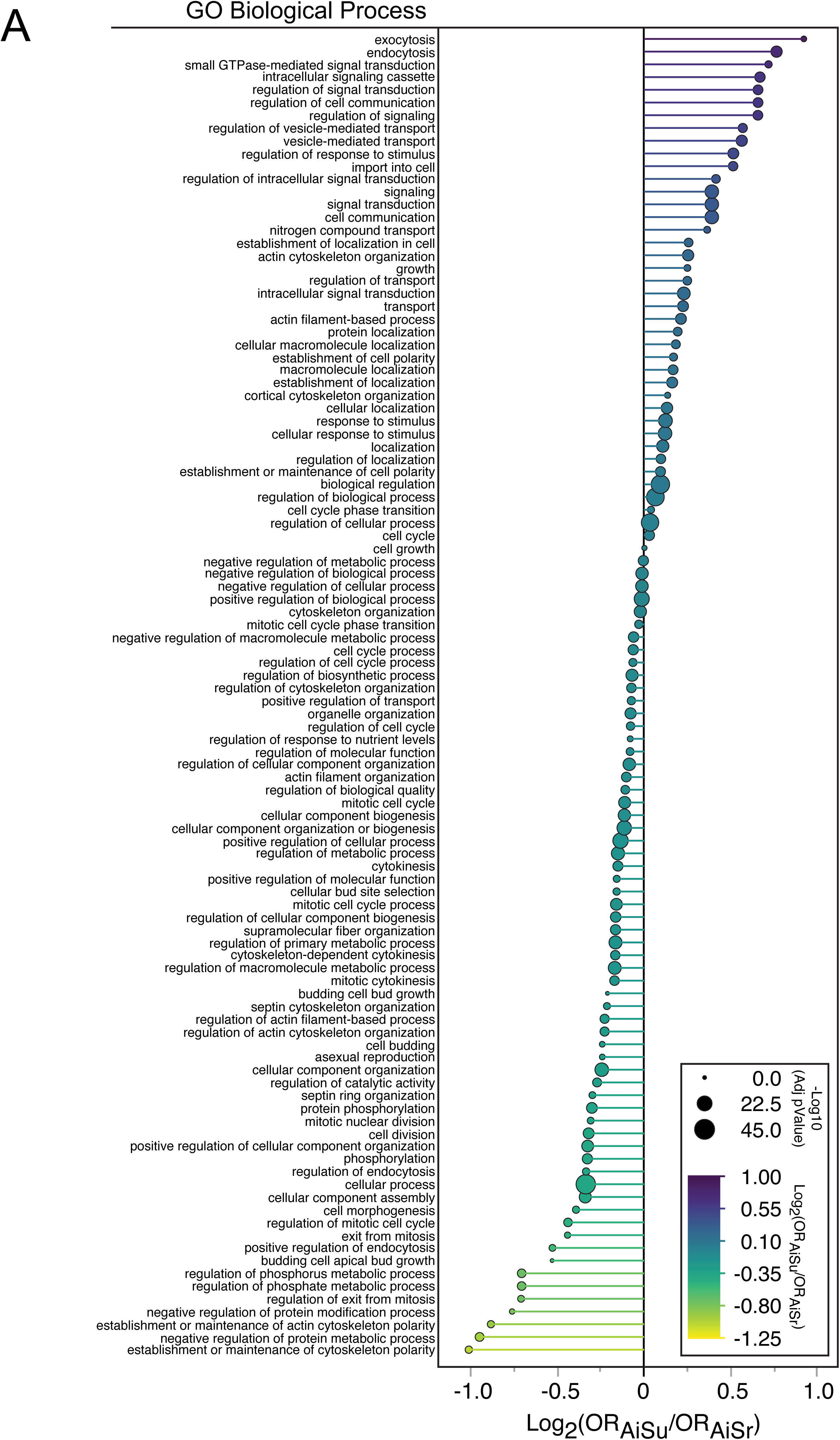
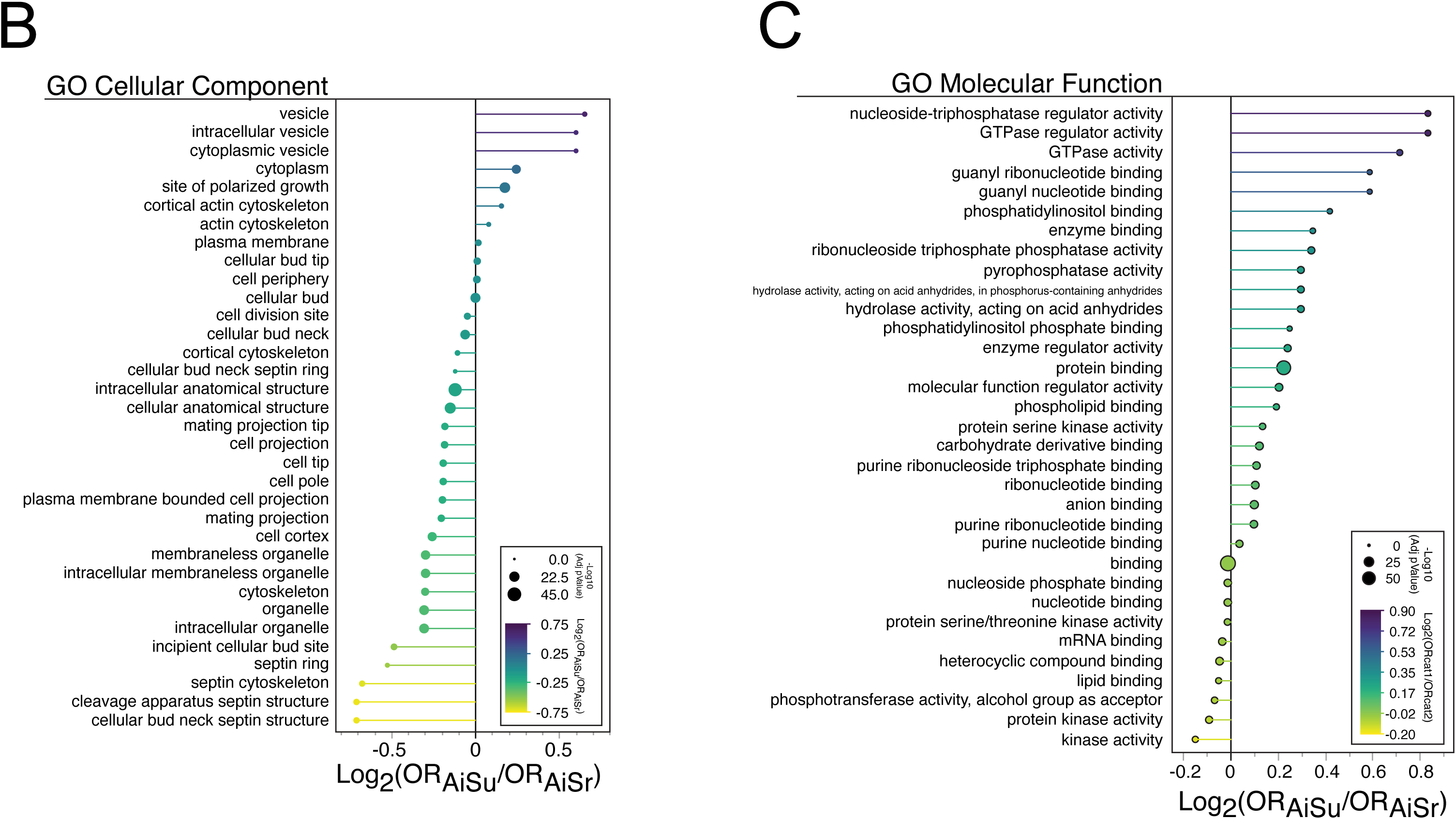
(A) Comprehensive comparison of log odds ratios for biological function GO terms enriched in AiSu versus AiSr response classes. (See B and C below). (B) Complete comparison of log odds ratios for cellular component GO terms enriched in AiSu versus AiSr response classes. (C) Complete comparison of log odds ratios for molecular function GO terms enriched in AiSu versus AiSr response classes.

**Supplemental Fig. S3.**
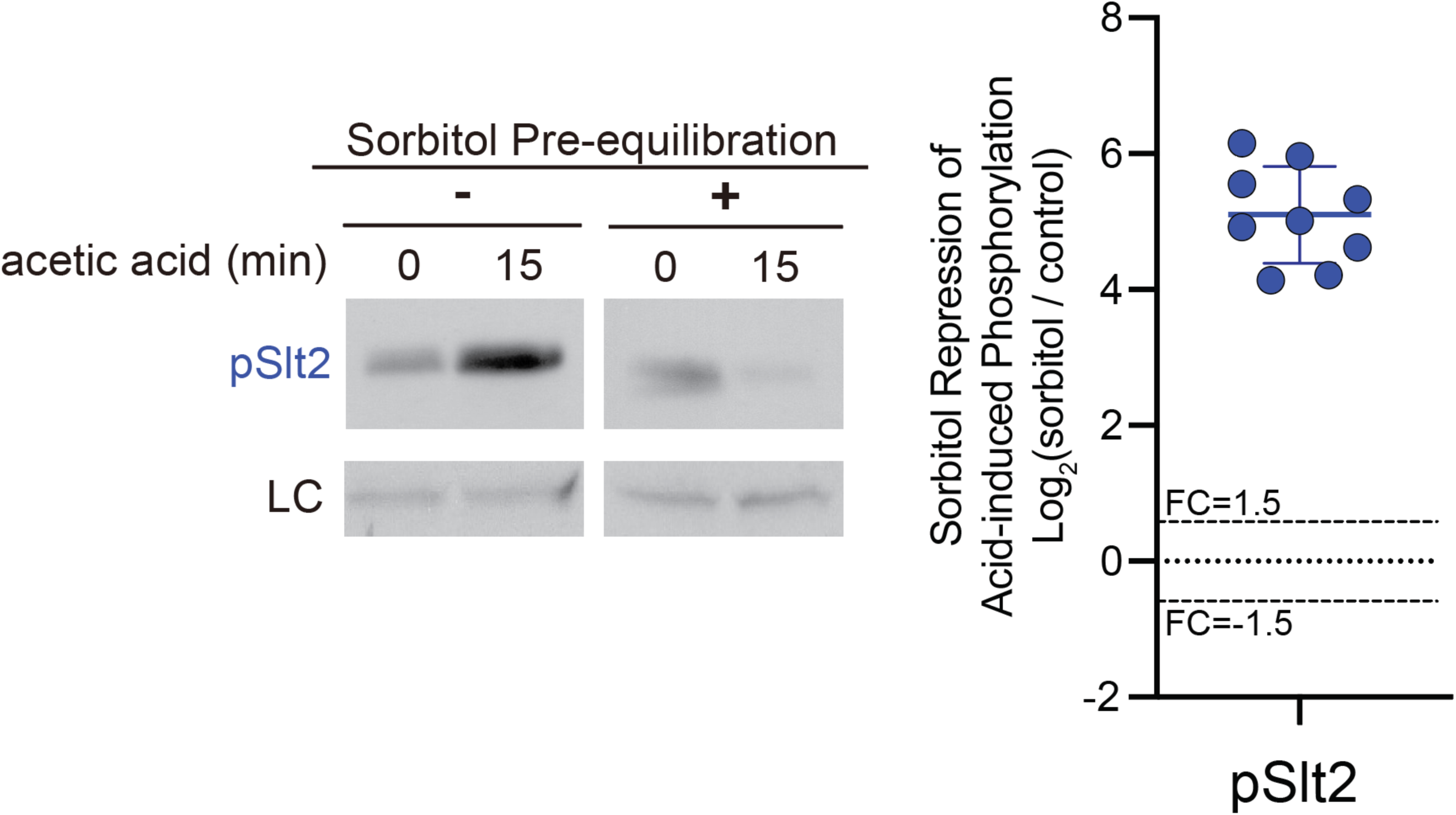
(Left) Immunoblots of phosphorylated Slt2 (pSlt2) and loading control (LC) in response to acid stress with or without sorbitol pre-equilibration. (right) Quantitative analysis of replicate pSlt2 immunoblots.

**Supplemental Fig. S4.**
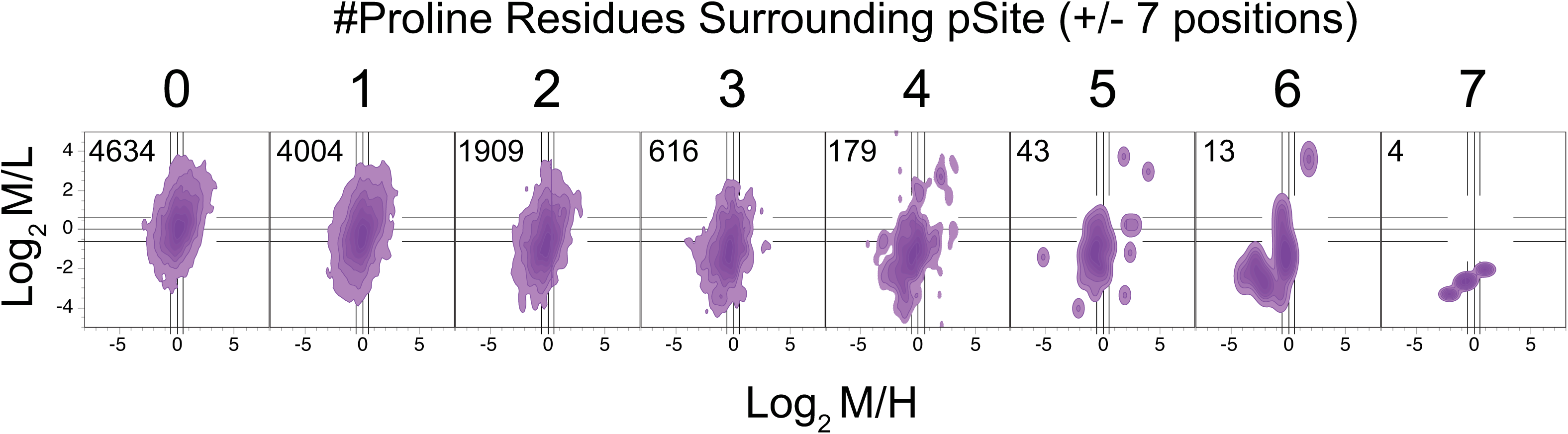
Distribution of proline-containing phosphosites (pSites) in the acid vs control (Log_2_M/L) and sorbitol-equilibrated acid vs control (Log_2_M/H) response space.

**Supplemental Fig. S5.**
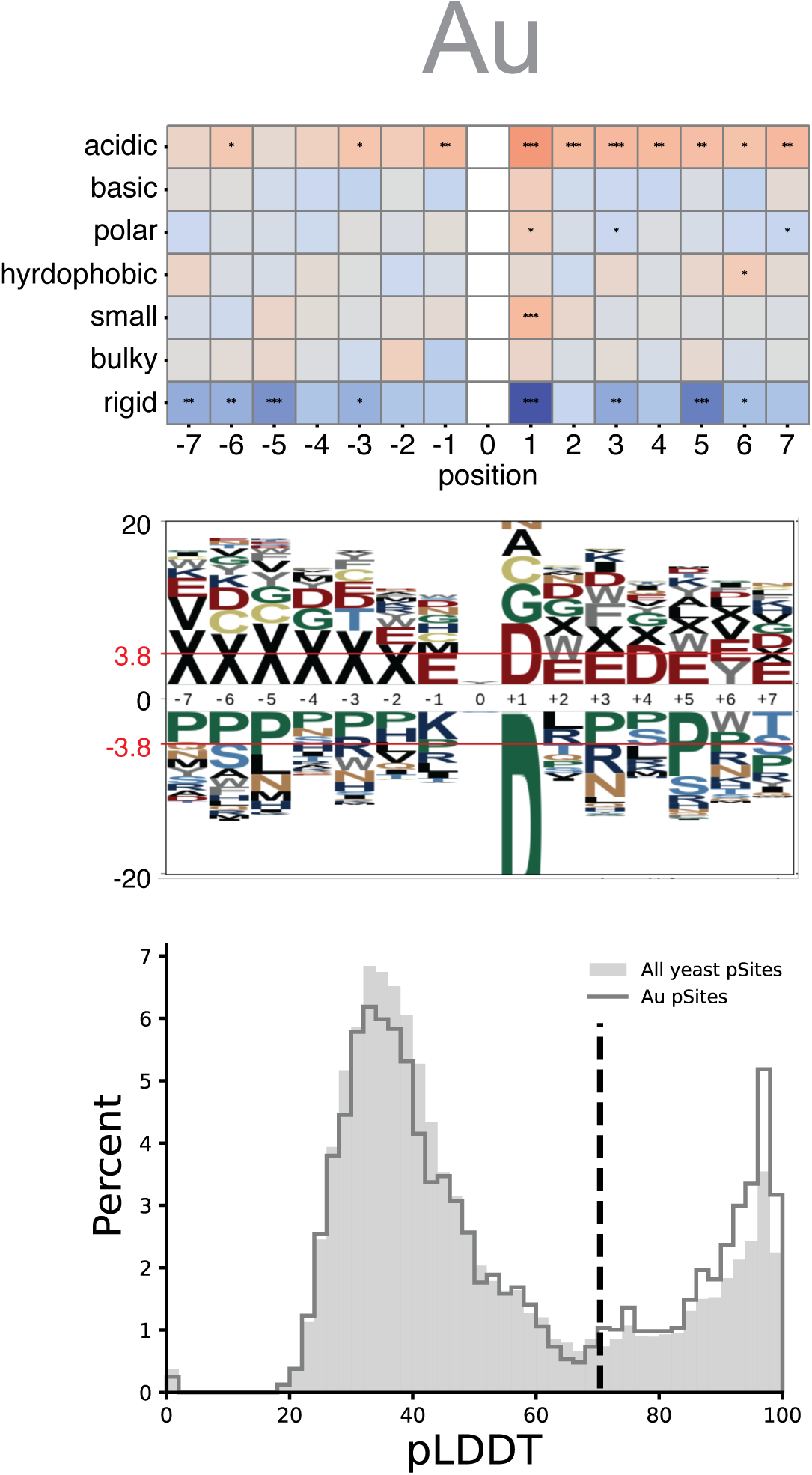
Analysis of Au response class motifs. (top) Position-specific enrichment of local physicochemical properties (top) and amino acids (middle) surrounding the acid responsive phosphosite (position 0). Significant enrichment is denoted by the red line with log odds > 3.8. pLogo plot is zoomed to visualize enriched residues that would be difficult to see fully unmagnified due to extreme under enrichment of proline at the +1 position. (bottom) Plot of AlphaFold 3 pLDDT score distributions in the Au response class (solid line) versus the distribution of all publicly curated experimental yeast phosphosites (shaded area). pLDDT scores >70 are accurately predicted fold structures (dashed grey line), while those <70 are low accuracy predictions that often coincide with less well-ordered structures. X represents gap filling letter indicating the absence of sequence. X’s to the left indicate that the sequences are from the N-terminal end of the protein whose phosphorylation sites is in the Au response class.

**Supplemental Fig. S6.**
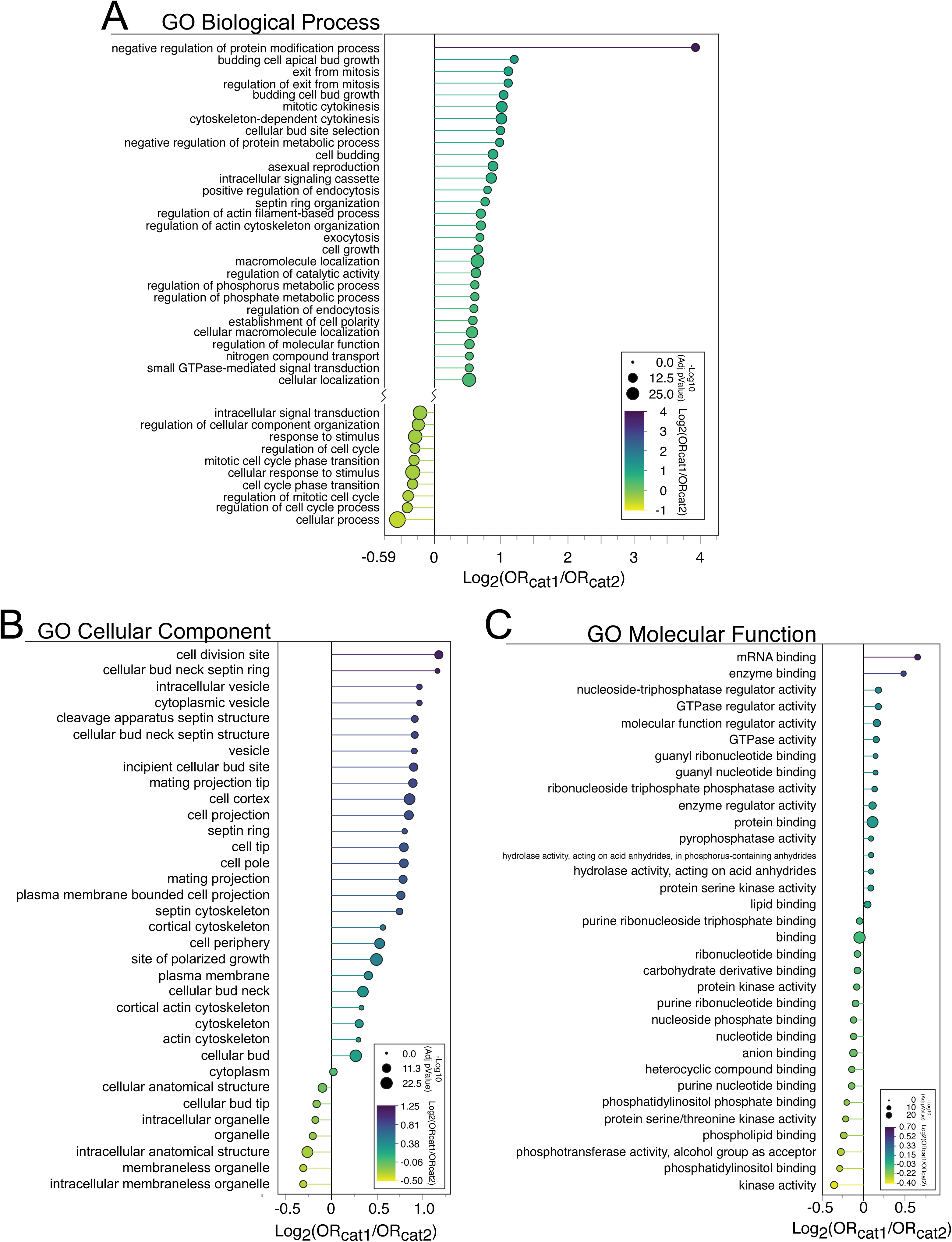
(A) Complete comparison of log odds ratios for biological function GO terms enriched in Cat1 versus Cat2 phosphosites. (B) Complete comparison of log odds ratios for cellular component GO terms enriched in Cat1 versus Cat2 phosphosites. (C) Complete comparison of log odds ratios for molecular function GO terms enriched in Cat1 versus Cat2 phosphosites.

**Supplemental Fig. S7.**
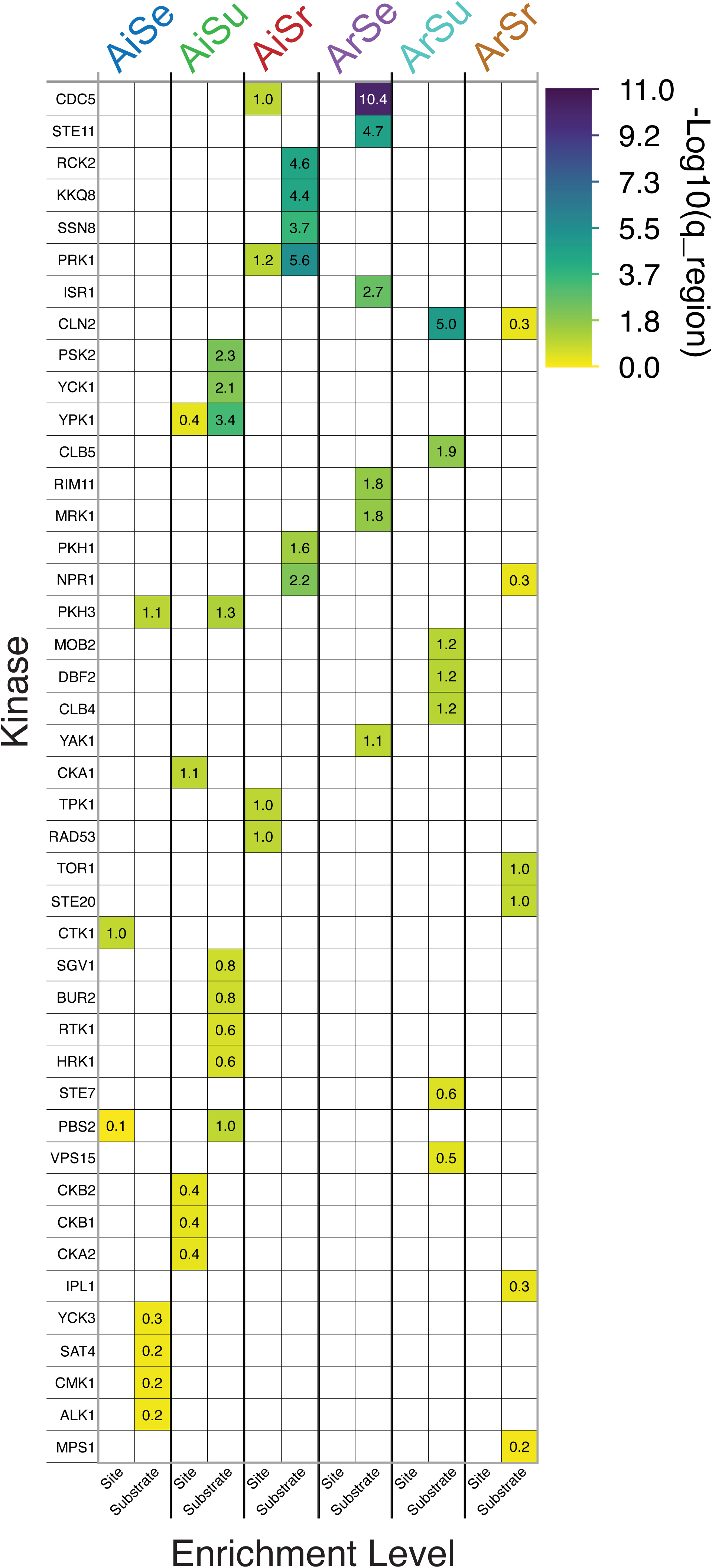
Complete result for kinase-site and kinase-substrate enrichment in the 6 major acid response classes.

**Supplemental Fig. S8.**
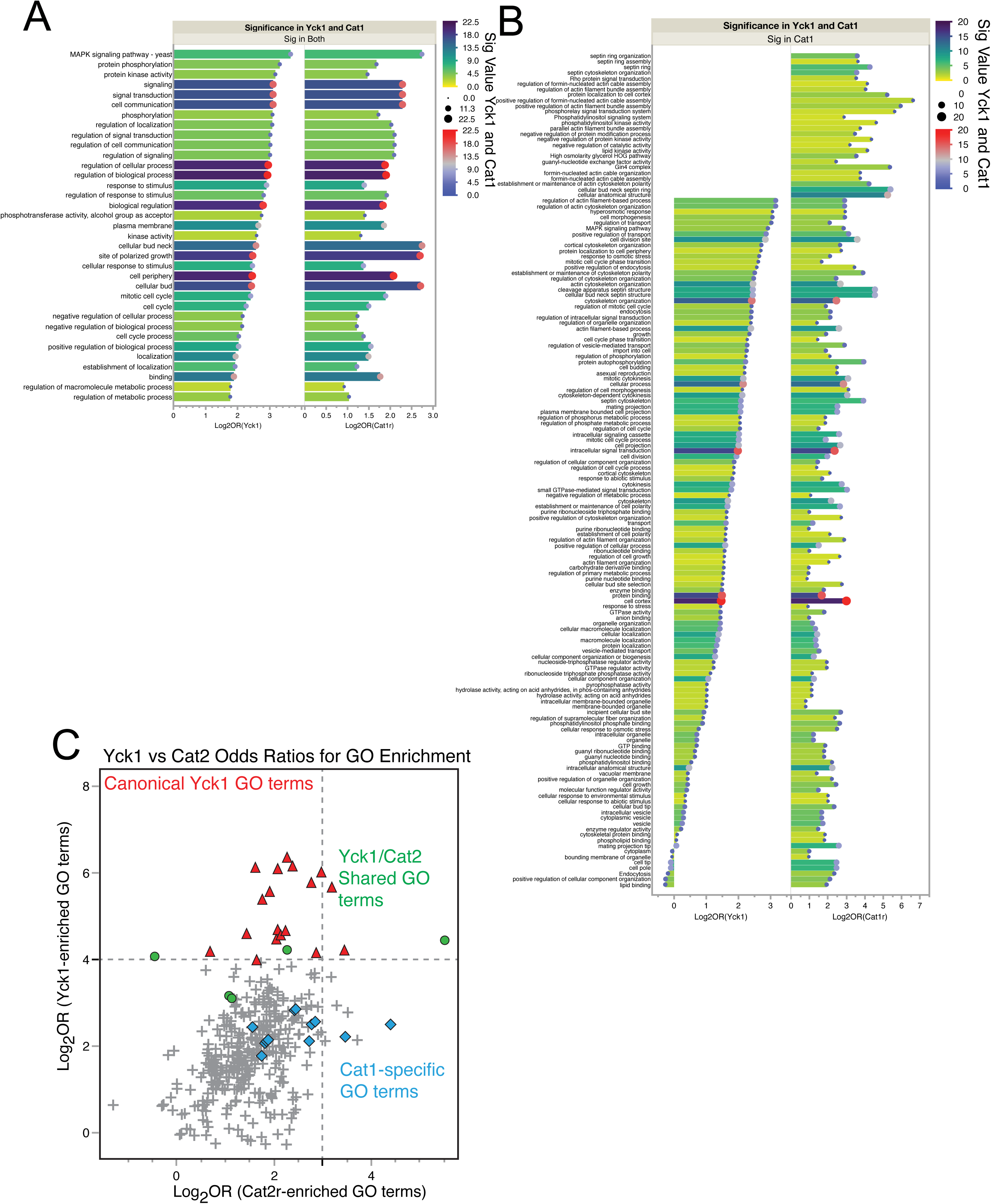
(A) Complete comparison of Yck1-specific versus Cat1-specific GO terms found to be significant for both cases. (B) Comparison of Yck1-specific versus Cat1-specific GO terms filtered for terms that are significant for Cat1 phosphosites, specifically. (C) Correlation plot of log odds ratios for GO terms enriched by Yck1 versus Cat2 phosphosites (basophilic sites common to the AiSr response class). Cat1-enriched GO terms serve as the color metric to show that Cat2 phosphosites do not equivalently enrich the same GO terms as Cat1 phosphosites. Cat1r, Cat1 phosphosites restricted to the boundaries: Log_2_M/L >0, - 1<Log_2_M/H<1. Cat2r, Cat2 phosphosites restricted to the boundaries: Log_2_M/L >0, Log_2_M/H >1.

**Supplemental Fig. S9.**
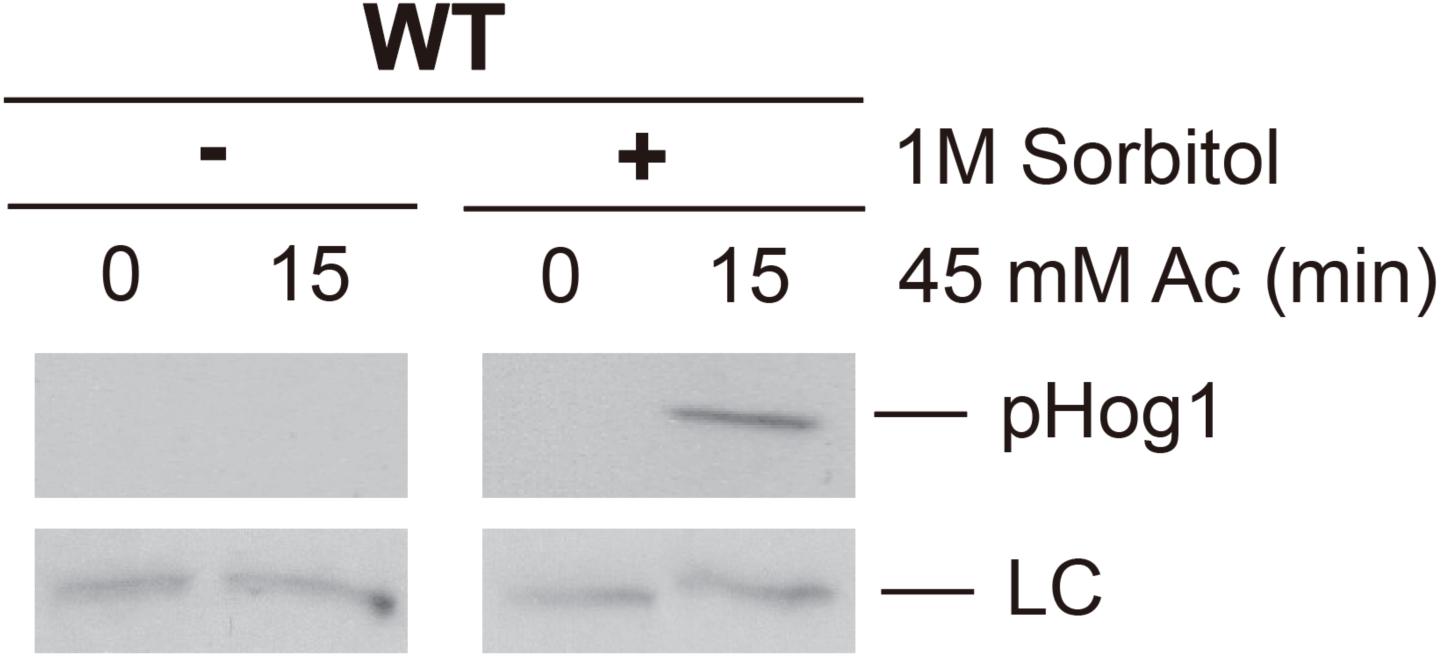
Immunoblot of phosphorylated/activated Hog1 map kinase in cells stressed with acetic acid with and without sorbitol pre-equilibration. LC, loading control.

