## Supplementary material for "Deep quantitative phosphoproteomics identifies non-canonical pH-sensitive yeast phosphorylation networks": Su et al 2026_SUPPLEMENTAL

Data within this document include:

**Supplemental Fig. S1 –** Volcano plots for M/H and M/L responses
**Supplemental Fig. S2 –** Gene ontology enrichments for the major acid response classes
**Supplemental Fig. S3 –** Quantitative immunoblotting for phospho-Slt2 in acid stress with and without sorbitol pre-equilibration
**Supplemental Fig. S4 –** Distribution of proline-containing phosphosites (pSites) in the acid stress response
**Supplemental Fig. S5 –** Analysis of Au response class phosphosite sequences and AlphaFold 3 structural predictions
**Supplemental Fig. S6 –** g:Profiler GO enrichment comparisons between Cat1 and Cat2 phosphosites
**Supplemental Fig. S7 –** Complete results for kinase-site and kinase-substrate enrichments across the 6 major acid response classes.
**Supplemental Fig. S8 –** Complete comparison of GO terms enriched by Yck1 versus Cat1r protein networks. Comparison of Yck1 and Cat2r protein networks
**Supplemental Fig. S9 –** Immunoblot for phospho-Hog1 in acid stress with and without sorbitol pre-equilibration

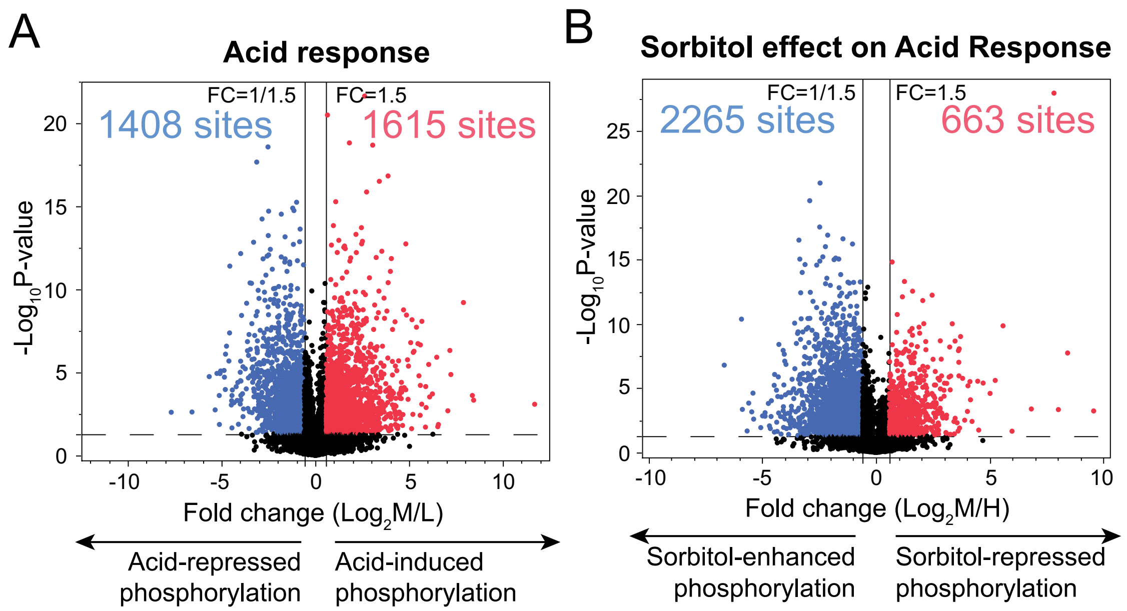


**Supplemental Fig. S1.** (A) Volcano plot for acid vs control ($Log_2M/L$). (B) Volcano plot for sorbitol-equilibrated acid vs control ($Log_2M/H$). (Dashed horizontal line) Significance threshold $p = 0.05$. (Solid vertical lines) $Log_2$ responses representing a -1.5 and 1.5-fold change.

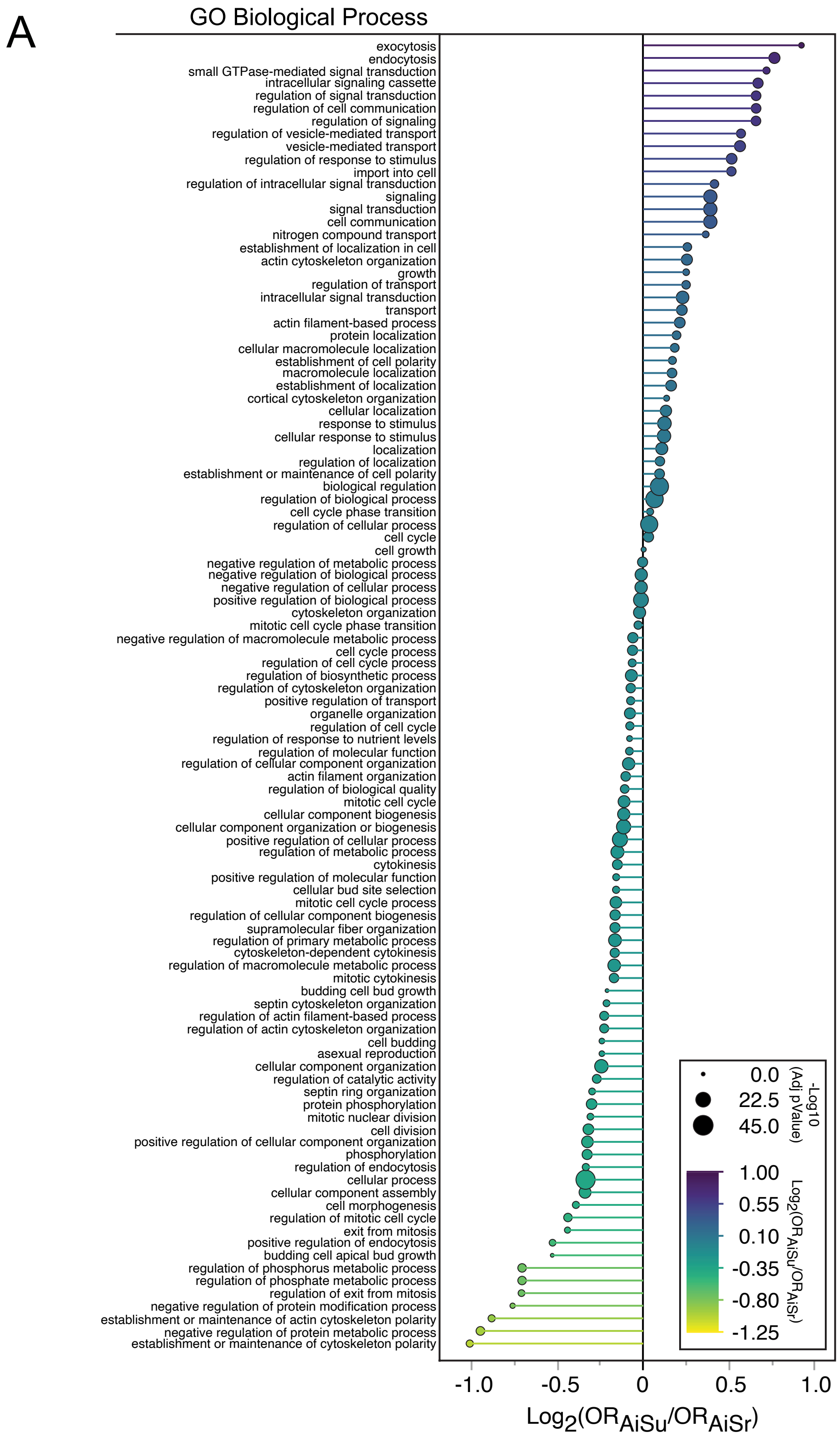


**Supplemental Fig. S2.** (A) Comprehensive comparison of log odds ratios for biological function GO terms enriched in AiSu versus AiSr response classes. (See B and C below).

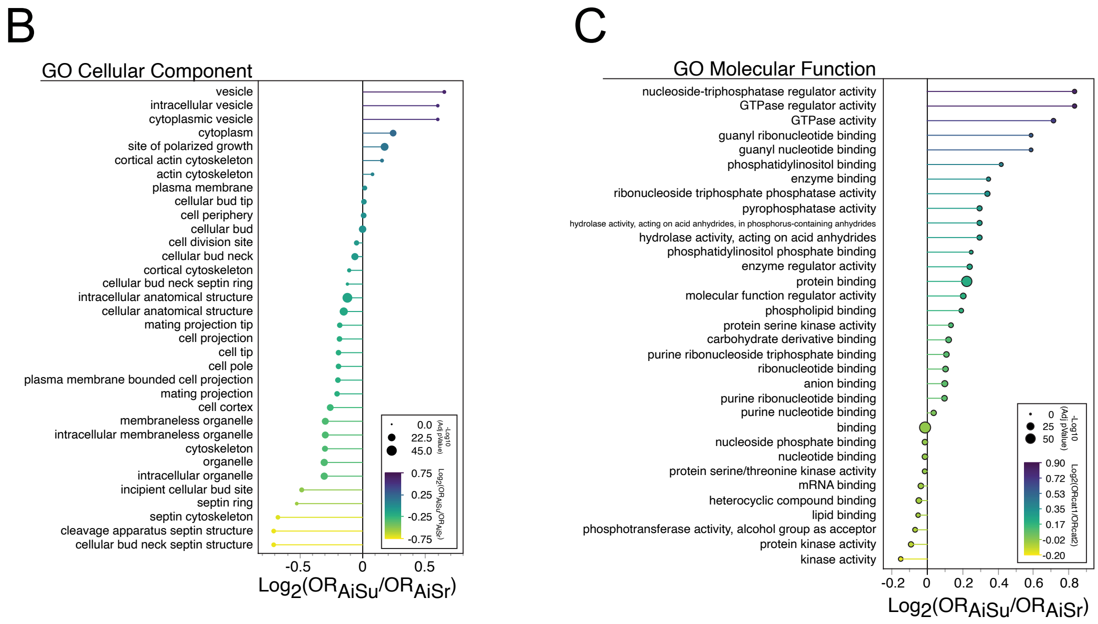


**Supplemental Fig. S2 (continued).** (B) Complete comparison of log odds ratios for cellular component GO terms enriched in AiSu versus AiSr response classes. (C) Complete comparison of log odds ratios for molecular function GO terms enriched in AiSu versus AiSr response classes.

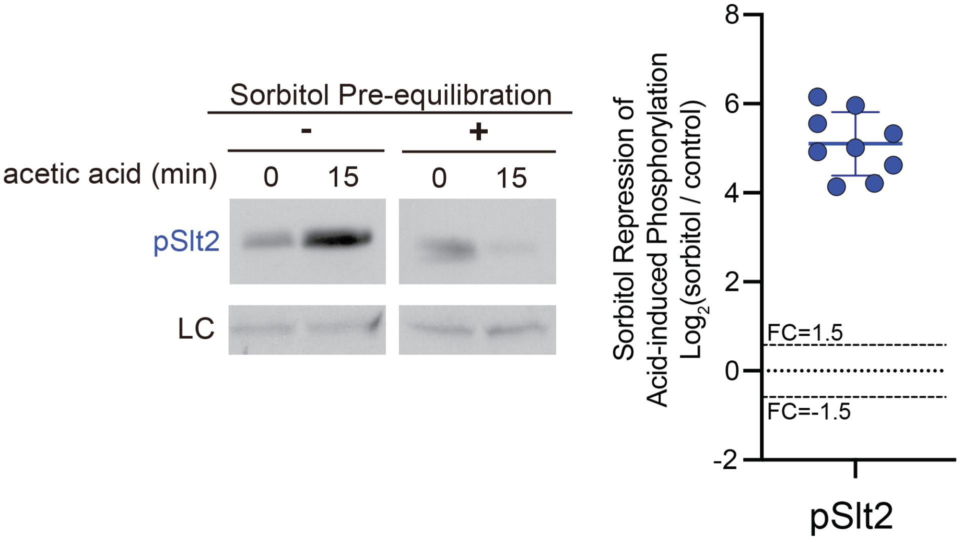


**Supplemental Fig. S3.** (Left) Immunoblots of phosphorylated Slt2 (pSlt2) and loading control (LC) in response to acid stress with or without sorbitol pre-equilibration. (right) Quantitative analysis of replicate pSlt2 immunoblots.

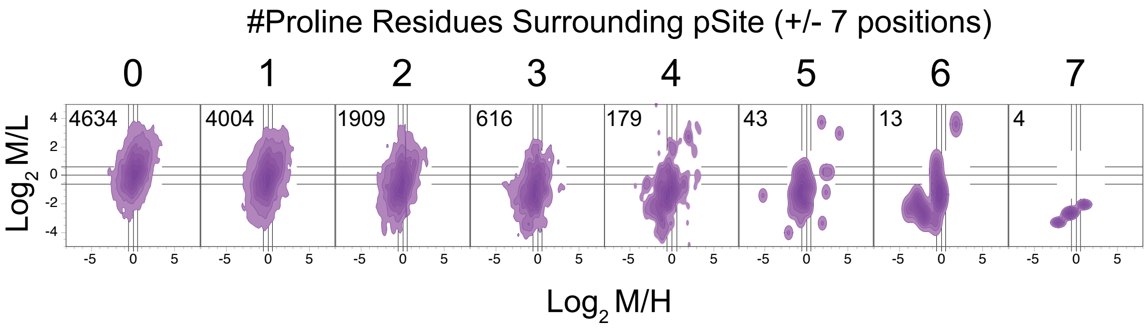


**Supplemental Fig. S4.** Distribution of proline-containing phosphosites (pSites) in the acid vs control ($Log_2M/L$) and sorbitol-equilibrated acid vs control ($Log_2M/H$) response space.

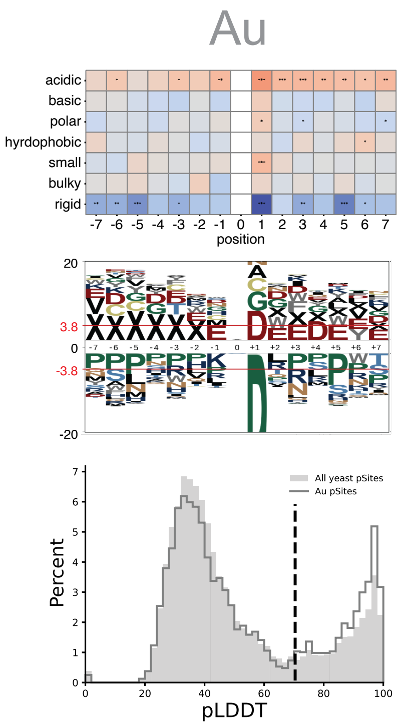


**Supplemental Fig. S5.** Analysis of Au response class motifs. (top) Position-specific enrichment of local physicochemical properties (top) and amino acids (middle) surrounding the acid responsive phosphosite (position 0). Significant enrichment is denoted by the red line with log odds > 3.8. pLogo plot is zoomed to visualize enriched residues that would be difficult to see fully unmagnified due to extreme under enrichment of proline at the +1 position. (bottom) Plot of AlphaFold 3 pLDDT score distributions in the Au response class (solid line) versus the distribution of all publicly curated experimental yeast phosphosites (shaded area). pLDDT scores >70 are accurately predicted fold structures (dashed grey line), while those <70 are low accuracy predictions that often coincide with less well-ordered structures. X represents gap filling letter indicating the absence of sequence. X's to the left indicate that the sequences are from the N-terminal end of the protein whose phosphorylation sites is in the Au response class.

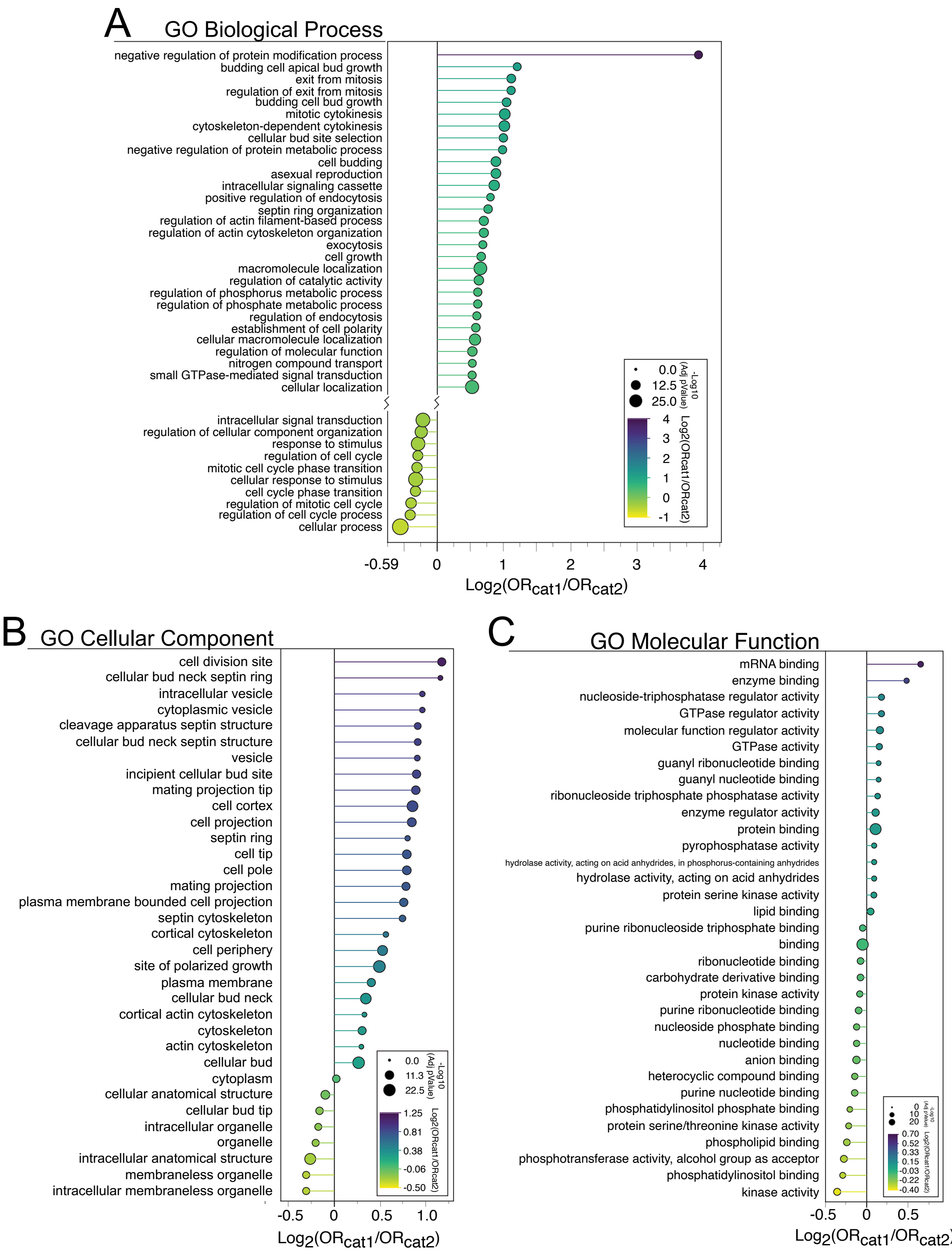


**Supplemental Fig. S6.** (A) Complete comparison of log odds ratios for biological function GO terms enriched in Cat1 versus Cat2 phosphosites. (B) Complete comparison of log odds ratios for cellular component GO terms enriched in Cat1 versus Cat2 phosphosites. (C) Complete comparison of log odds ratios for molecular function GO terms enriched in Cat1 versus Cat2 phosphosites.

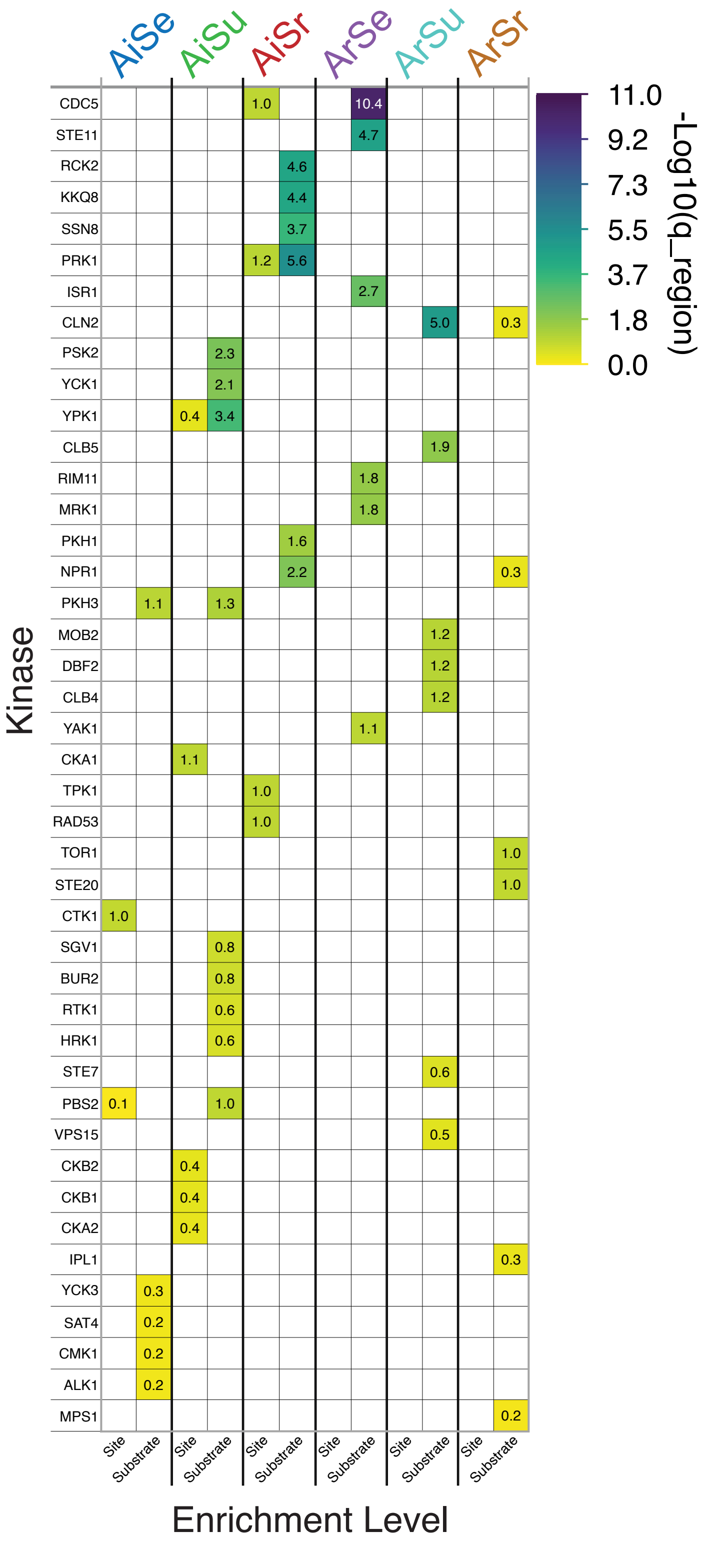


**Supplemental Fig. S7.** Complete result for kinase-site and kinase-substrate enrichment in the 6 major acid response classes.

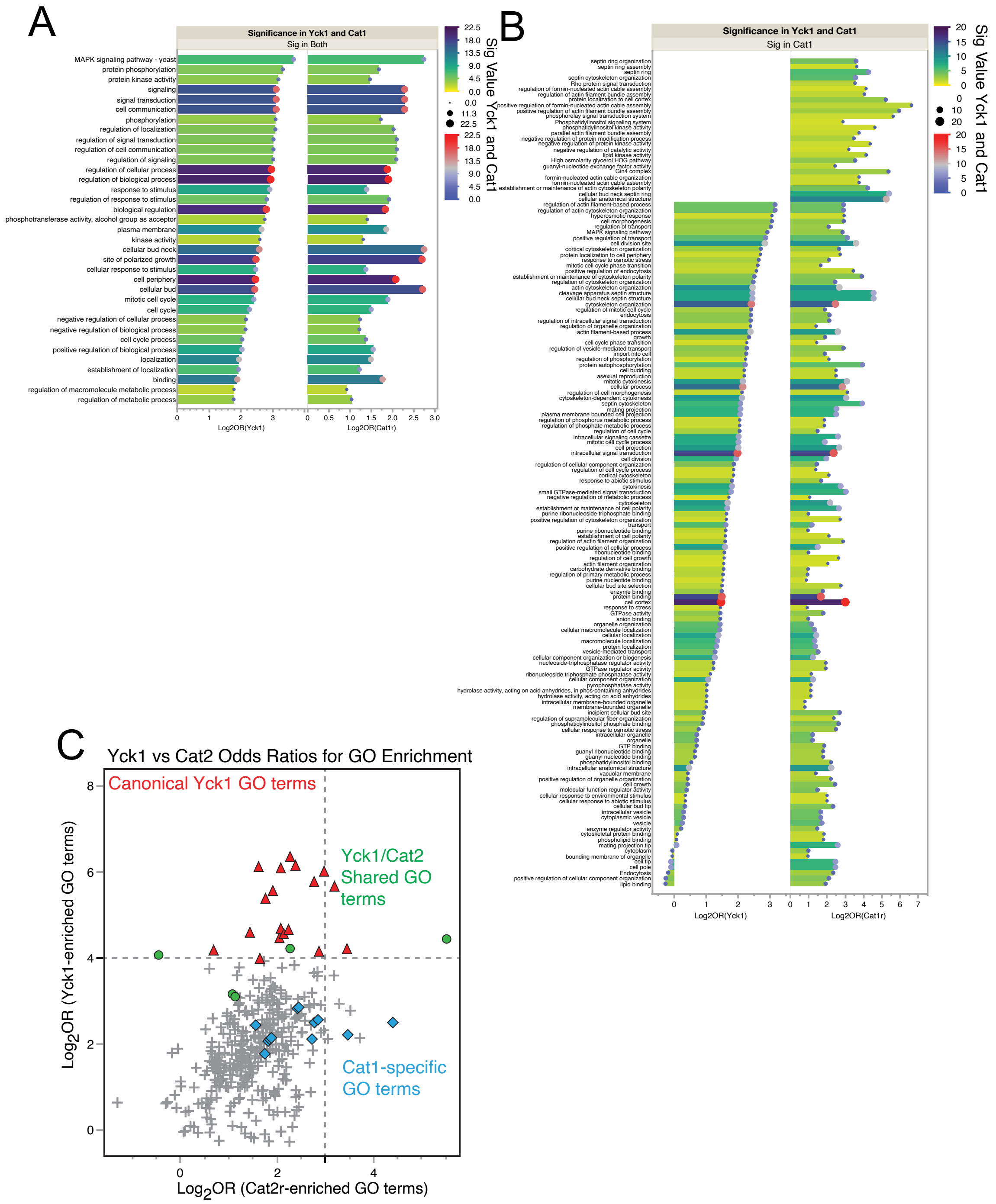


**Supplemental Fig. S8.** (A) Complete comparison of Yck1-specific versus Cat1-specific GO terms found to be significant for both cases. (B) Comparison of Yck1-specific versus Cat1-specific GO terms filtered for terms that are significant for Cat1 phosphosites, specifically. (C) Correlation plot of log odds ratios for GO terms enriched by Yck1 versus Cat2 phosphosites (basophilic sites common to the AiSr response class). Cat1-enriched GO terms serve as the color metric to show that Cat2 phosphosites do not equivalently enrich the same GO terms as Cat1 phosphosites. Cat1r, Cat1 phosphosites restricted to the boundaries: $Log_2M/L > 0$, $-1 < Log_2M/H < 1$. Cat2r, Cat2 phosphosites restricted to the boundaries: $Log_2M/L > 0$, $Log_2M/H > 1$.

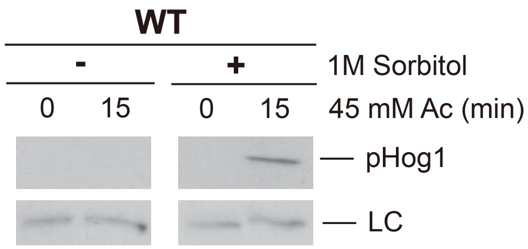


**Supplemental Fig. S9.** Immunoblot of phosphorylated/activated Hog1 map kinase in cells stressed with acetic acid with and without sorbitol pre-equilibration. LC, loading control.
